# Mosquito host background influences microbiome-ZIKV interactions in field and laboratory-reared *Aedes aegypti*

**DOI:** 10.1101/2025.02.02.636091

**Authors:** Cintia Cansado-Utrilla, Miguel A. Saldaña, George Golovko, Kamil Khanipov, Alex L. Wild, Laura E. Brettell, Scott C. Weaver, Eva Heinz, Grant L. Hughes

**Affiliations:** Departments of Vector Biology and Tropical Disease Biology, Liverpool School of Tropical Medicine, Liverpool, UK; Mosquito and Vector Control Division, Harris County Public Health, Texas, USA; Department of Microbiology and Immunology, Institute for Human Infections and Immunity, University of Texas Medical Branch, Galveston, Texas, USA; Department of Pharmacology, The University of Texas Medical Branch, Texas, USA; Department of Integrative Biology, University of Texas, Austin, Texas, United States of America; School of Science, Engineering and Environment, University of Salford, Manchester, M5 4WT, UK; Departments of Vector Biology and Clinical Sciences, Liverpool School of Tropical Medicine, Liverpool, UK; Department of Microbiology and Industrial Biotechnology, Institute of Pharmacy & Biomedical Sciences, University of Strathclyde, Glasgow, UK

## Abstract

The mosquito microbiota represents an intricate assemblage of microorganisms, comprising bacteria, fungi, viruses, and protozoa. Factors modulating microbiome abundance and composition include host genetic background, environmental parameters, and pathogen exposure. Conversely, the microbiome profoundly influences pathogen infection of the mosquito host and thus harbours considerable potential to impact the transmission of vector-borne diseases. As such, there is a growing interest in using the microbiome in novel vector-control strategies, including exploiting the natural ability of some microbes to interfere with infection of the vectors by pathogens. However, before novel microbiome-based vector control approaches can move towards translation, a more complete understanding of the interactions between mosquitoes, their microbiome, and the pathogens they transmit, is required to better appreciate how variation in the microbiome of field mosquitoes affects these interactions. To examine the impact of the host background and the associated diversity of microbiomes within distinct hosts, but without artificially manipulating the microbiome, we exposed several laboratory-reared and field-collected *Aedes aegypti* mosquito lines to Zika virus (ZIKV) and correlated their microbial load and composition to pathogen exposure and viral infection success. We observed significant differences in ZIKV exposure outcomes between the different mosquito lines and their associated microbiomes, and found that ZIKV alteration of the microbiomes was distinct in different lines. We also identified microbial taxa correlating with either ZIKV infection or a lack of infection. In summary, our study provides novel insights into the variability of pathogen interactions within the mosquito holobiont. A more complete understanding of which factors influence the tripartite interactions between *Aedes* mosquitoes, their microbiome, and arboviral pathogens, will be critical for the development of microbial-based interventions aimed at reducing vector-borne disease burden.

**Author summary.:** The mosquito microbiome composition differs within an individual across its development, as well as between individual mosquitoes at the same developmental stage, and between spatially or genomically different mosquito populations. The microbiome is highly relevant for the ability of mosquitoes to transmit pathogens. Furthermore, certain microbes have been shown to influence pathogen infection of the mosquito, while conversely, infection with a pathogen can alter the mosquito microbiome. However, we have a poor understanding how universally conserved these pathogen-related effects observed in a specific host-microbiome combination are in different mosquito populations with their respective microbiomes. To address this, we infected different mosquito lines, either reared in the laboratory or caught in the field and examined the microbiomes after exposure to Zika virus (ZIKV) compared to unchallenged microbiomes. We also examined how the virus infection progressed in different mosquito lines and correlations with further microbiome changes. The observed microbiome responses differed between host lines, potentially due to either different microbiomes associated with the respective hosts. Alternatively, the host may respond differently to the viral infection, which subsequently alters the microbiome in a distinct manner, or a combination of host and microbiome effects may occur. As microbes are being evaluated for novel approaches to control mosquito-borne disease, our findings are highly relevant to contribute to a more complete understanding of host-microbe interactions which will be critical to develop these approaches. Variation of the microbiome of different mosquito lines need to be considered in experimental designs and when interpreting results from specific studies. It is especially relevant for deployment of interventions in the field where microbial variability is known to be higher and where variation is observed between mosquito populations.

## Background

The mosquito and its associated microbial community collectively form the mosquito holobiont, a complex ecosystem with multi-layered interactions [1]. The host-microbe interactions influence several phenotypes of the mosquito host such as growth and development, reproduction, and the ability to transmit pathogens, all of which are important for vectorial capacity [2]. The microbiome composition is influenced by the mosquito host genetic background but also multiple other factors including environmental parameters, microbe-microbe interactions and exposure to pathogens [3–9]. Variability of microbiomes could therefore be an explanation for the variation seen in the vector competence of different mosquito lines of the same species [10–14].

Interactions between microbes and pathogens are bi-directional and include direct and indirect effects, with the microbiome affecting the outcomes of infection with human pathogens, and conversely pathogen infection altering the microbiome composition and abundance. Bi-directional interactions can be mediated by insect immunity, given that both pathogens and microbes elicit and are modulated by these pathways [15, 16]. Additionally, microbes can directly affect pathogen infection via the production of compounds affecting the parasites or arboviruses [17–19]. These direct microbiome-pathogen interactions can both positively and negatively affect mosquito susceptibility to pathogens. For instance, in *Aedes aegypti,* some isolates of *Serratia* have been implicated in enhancing susceptibility to dengue virus (DENV) infection, whereas members of the *Rosenbergiella* genus impair vector competence to both DENV and Zika virus (ZIKV) [17, 19]. Whilst these studies focus on specific bacterial taxa with distinct effects in particular host lines, we were interested in understanding how the collective microbiome interacts with arboviruses and vice versa, and how conserved the observed interactions are between different host backgrounds.

In addition, much of our insight on the tripartite interactions between the host, their microbes, and pathogens, is derived from laboratory-based studies on long-term, inbred mosquito lines, where the involvement of the microbiome is often assessed by perturbation. This is typically achieved by administration of antibiotics to alter the microbiome; however, this approach also impacts host fitness and mitochondria. It does not necessarily completely clear the microbiota, but rather generates a highly artificial situation of a limited or a heavily biased microbiome [7, 20, 21]. Alternatively, microbes can be introduced into mosquitoes either at the aquatic stages in the larval water, or to adults via a sugar meal, and can thus be added to an already existing microbiome. This may reduce the level of disruption of the holobiont system, and mimick administration approaches that could occur in control interventions. Using this approach, field collected bacterial strains have been shown to modulate vector competence [6]. While such manipulation experiments provide evidence for the microbiomes’ role in vector competence, and in the case of the latter, provide candidates for microbial control, they do not comprehensively address how variability in the microbiome influences tripartite interactions.

Exploiting the natural microbiome variation observed in mosquitoes, and particularly those in the field, offers a potential avenue to further explore the role of the microbiome on mosquito phenotypes, including vector competence. In this study, we used this natural microbiome variability to examine tripartite interactions between distinct *Ae. aegypti* mosquito lines, their microbiomes, and ZIKV. To address how differences in the microbiota between and within mosquito populations altered interactions with ZIKV, we collected host-seeking females from different geographic regions, provided them with an infectious ZIKV blood meal, and monitored viral infection status, viral loads post infection, and microbiome composition. Additionally, using two different laboratory-reared *Ae. aegypti* colonies, we examined if microbiomes responded to pathogen infection in a similar fashion in differing host backgrounds. We show that different mosquito lines, that have difference in host genetics and associated microbiomes, can profoundly alter ZIKV-microbiome interactions. Our results highlight the complexity of tripartite interactions in mosquitoes, and are important to consider for the development of microbial-based control strategies.

## Methods

### Mosquito lines

Field mosquitoes were collected outdoors over a three-day period, in Austin, Galveston, and Brownsville (Texas, USA). On each day, host-seeking mosquitoes were captured using CDC Fay-Prince traps for three hours at dawn and dusk, with collection cups replaced every hour. Mosquitoes were retrieved from traps and stored in large cartons kept within plastic bins containing a moist sponge for humidity and provided with 10% sucrose until their arrival at the insectaries of the University of Texas Medical Branch (UTMB) (Galveston, Texas, USA). Mosquitoes were then anesthetized at 4°C and their species and sex were determined by morphological identification. Female *Ae. aegypti* were transferred to new cages. Laboratory reared mosquito lines used in this study were Galveston and Rio Grande Valley (RGV), two recently established colonies at UTMB, the former for three generations and the latter for six. All mosquito lines were maintained under standard insectary conditions at UTMB (27°C and 80% humidity) and fed with 10% sucrose.

### Viral strains and mosquito infections

The viral strain used in this study was ZIKV MEX 1-7 (KX247632.1), isolated from *Ae. aegypti* in Mexico in 2016 [22]. The virus was acquired as a lyophilized stock from the World Reference Center for Emerging Viruses and Arboviruses at UTMB. It was cultured in C6/36 cells, an *Ae. albopictus*-derived cell line, followed by four passages in the mammalian Vero cell line to generate stocks. Vero cells were maintained in high-glucose Dulbecco’s modified Eagle’s medium (DMEM) supplemented with 5% foetal bovine serum (FBS) and 1% penicillin/streptomycin at 37°C and 5% CO_2_. Cages of laboratory-reared and field-collected mosquitoes were starved for 18 hours before being offered a blood meal spiked with ZIKV (10^6^ FFU/ml) (Austin N=113, Galveston N=40, Brownsville N=19, Galveston-lab N=57, RGV-lab N=85). Bloodmeals were offered five days post-pupal eclosion to lab mosquitoes and one to three days post collection to field mosquitoes. Mosquitoes that did not feed were removed. Galveston and RGV lab-reared mosquitoes were offered an uninfected bloodmeal (Galveston-lab N=40, RGV-lab N=40) as a control. Ten days after blood feeding, mosquitoes were euthanised and assessed for ZIKV infection using focus forming assays, and the microbiome was characterised using qPCR and 16S rRNA amplicon sequencing (**Figure 1**).

**Figure 1.**
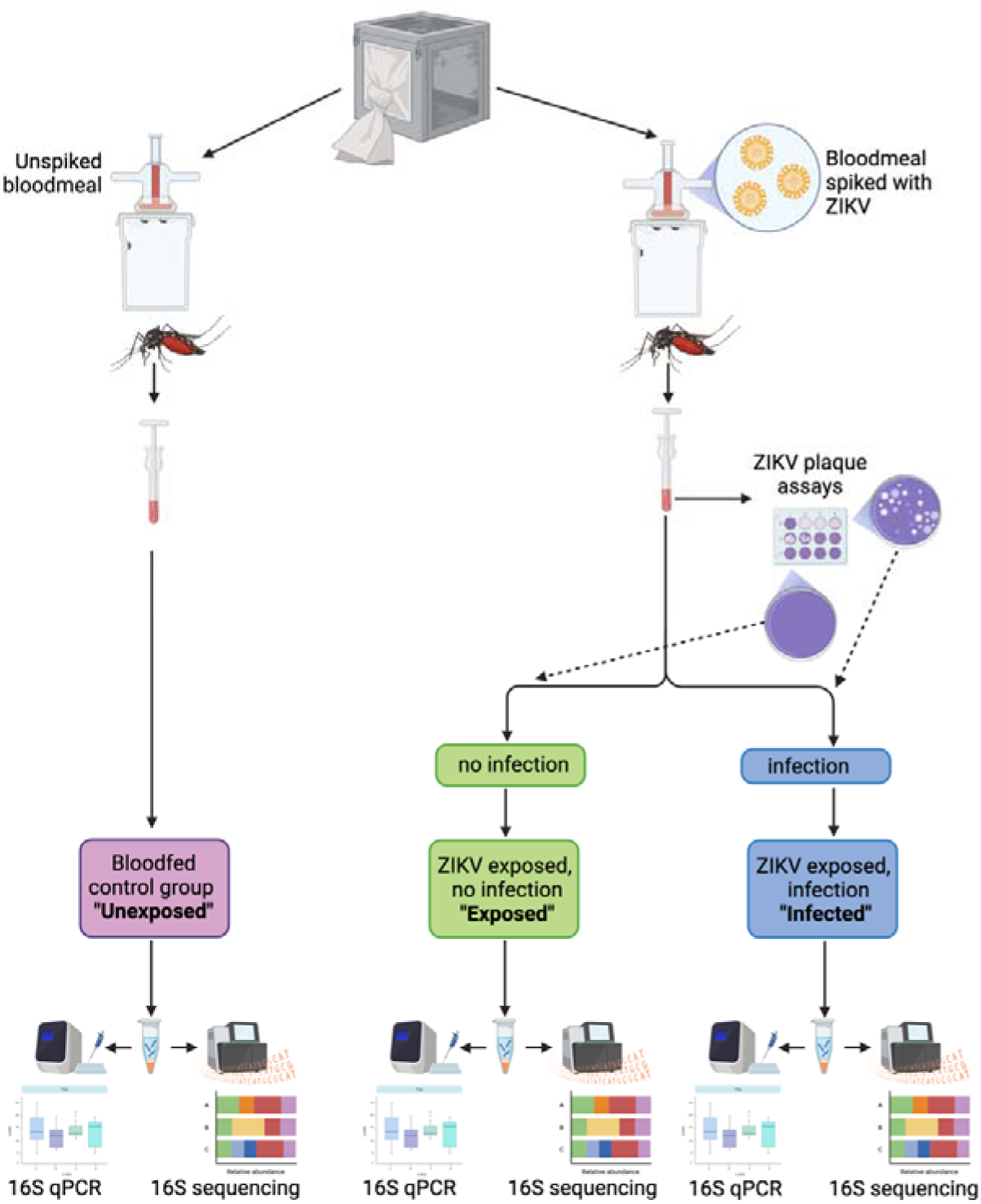
Experimental design for ZIKV infection of lab-reared *Ae. aegypti* lines. After ZIKV infectious blood meals mosquitoes were designated into groups termed “Exposed” indicating exposure but a lack of infection, or “Infected”, indicating infection of ZIKV in mosquitoes. An “Unexposed” group consisted of blood meal without virus.

### Focus forming assay

Individual mosquitoes that had fed on an infected bloodmeal were surface sterilized (5 minutes in 70% ethanol followed by three washes in PBS for five minutes each) and homogenized in 500 µl of tissue culture medium (DMEM supplemented with 5% FBS, 1% penicillin/streptomycin and 1% amphotericin) using a TissueLyser II (Qiagen) for five minutes at 60 Hz. Mosquito samples were serially diluted and inoculated onto Vero cells in 48-well plates and overlaid with 0.8% methylcellulose in DMEM. Mosquito bodies and legs were used to determine viral infection ro dissemination, respectively. Plates were washed with PBS, incubated at 37°C for four days and fixed with 50:50 methanol:acetone. Foci were stained using a mouse hyperimmune polyclonal anti-ZIKV primary antibody (World Reference Center for Emerging Viruses and Arboviruses, UTMB) and HRP-labelled goat anti-mouse secondary antibody (KPL, Gaithersburg, MD). ZIKV foci were then visualized using an aminoethylcarbazole (AEC) detection kit (Enzo Diagnostics, Farmingdale, NY) according to the manufacturer’s protocol.

### Estimation of bacterial density

Genomic DNA was extracted from 250 µl of the homogenate, obtained from the material used for focus forming assay, using the NucleoSpin Tissue kit (Macherey-Nagel) as previously described and used as template for qPCR [23]. Universal bacterial 16S rRNA primers and the housekeeping S7 gene primers were used as previously described [23–25]. Relative gene expression was calculated using the 2^-ΔΔCt^ method [26]. Microbiome load (16S/S7) data were analysed in RStudio (version 1.4.1717), density and Q-Q plots with the *ggpubr* package (version 0.6.0) and Shapiro-Wilk tests using the *stats* package (version 4.3.2) [27, 28]. The data was not normally distributed in any of the groups, so Wilcoxon-Rank Test was used to compare the means using the *ggpubr* package.

### Analysis of 16S rRNA amplicon sequences

Genomic DNA from all mosquitoes was then used for high-throughput sequencing targeting the bacterial 16S ribosomal RNA gene. Sequencing libraries for each isolate were generated using universal 16S rRNA V3-V4 region primers following Illumina 16S rRNA metagenomic sequencing library protocols [29]. The samples were barcoded for multiplexing using Nextera XT Index Kit v2. Sequencing was performed on an Illumina MiSeq instrument using a MiSeq Reagent Kit v2 (500 cycles). Quality control and taxonomical assignment of the resulting reads was performed using CLC Genomics Workbench 8.0.1 Microbial Genomics Module (http://www.clcbio.com). Low quality reads containing nucleotides with a quality threshold below 0.05 (using the modified Richard Mott algorithm), as well as reads with two or more unknown nucleotides or sequencing adapters were removed. Reference based OTU selection was performed using the SILVA SSU v128 97% database [30]. Sequencing of 16S failed for seven samples (five field collected individuals (Austin) and two unexposed individuals (RGV)). Chimeras were removed from the dataset if the absolute crossover cost was 3 using a k-mer size of 6. Data were then transferred to RStudio (version 1.4.1717) for subsequent analyses. Samples with fewer than 2,000 reads were removed (18 from Austin, one from Galveston-field, one from Brownsville, two from Galveston-lab and six from RGV-lab), resulting in a final data set comprising 359 samples (90 from Austin, 39 from Galveston-field, 18 from Brownsville, 95 from Galveston-lab and 117 from RGV-lab; (**Table S1; Figure S1)**). Data were then converted to a phyloseq object using the *Phyloseq* package [31]. Diversity parameters (Shannon entropy and Bray-Curtis distance) were assessed using the *vegan* package [32]. Shannon diversity index data were tested for normality using density and Q-Q plots and Shapiro-Wilk tests. All data groups failed tests for normality, so a Wilcoxon-Rank Test was used to compare the means. Overall differences in beta diversity between groups was carried out using permutational multivariate analysis of variance (PERMANOVA) testing using the ‘Adonis2’ function in the *vegan* package with subsequent pairwise testing using the *PairwiseAdonis* package [33]. Beta diversity was visualised using NMDS plots and ellipses were added to the plots using the ‘stat_ellipse’ function in *ggplot2* using the default 95% confidence levels assuming multivariate t-distribution [34]. Determination of differentially abundant taxa between groups was calculated using Analysis of compositions of microbiomes with bias correction (ANCOM-BC) [35]. A heatmap showing differentially abundant taxa in RGV-lab mosquitoes to Galveston-lab mosquitoes for each of the three groups (unexposed, exposed and infected) was generated using the *pheatmap* package using the ANCOM-BC results [36].

## Results

### Mosquito line influences the ZIKV-microbiome interaction

To investigate whether interactions between ZIKV and the microbiome differ when using *Ae. aegypti* from different backgrounds, we fed two laboratory-reared *Ae. aegypti* lines (Galveston-lab and RGV-lab) with either a non-infectious bloodmeal (unexposed control group) or a bloodmeal spiked with ZIKV. Subsequently, we assessed the latter group for viral infection and categorised them as exposed (no ZIKV infection detected) or infected (ZIKV infection detected in the midgut). Only a subset of mosquitoes developed an infection, and this percentage differed significantly between lines, with 44% infection in RGV-lab mosquitoes and 26% infection in Galveston-lab mosquitoes (Chi-square, *p*=0.04) (**Figure 2A**).

**Figure 2.**
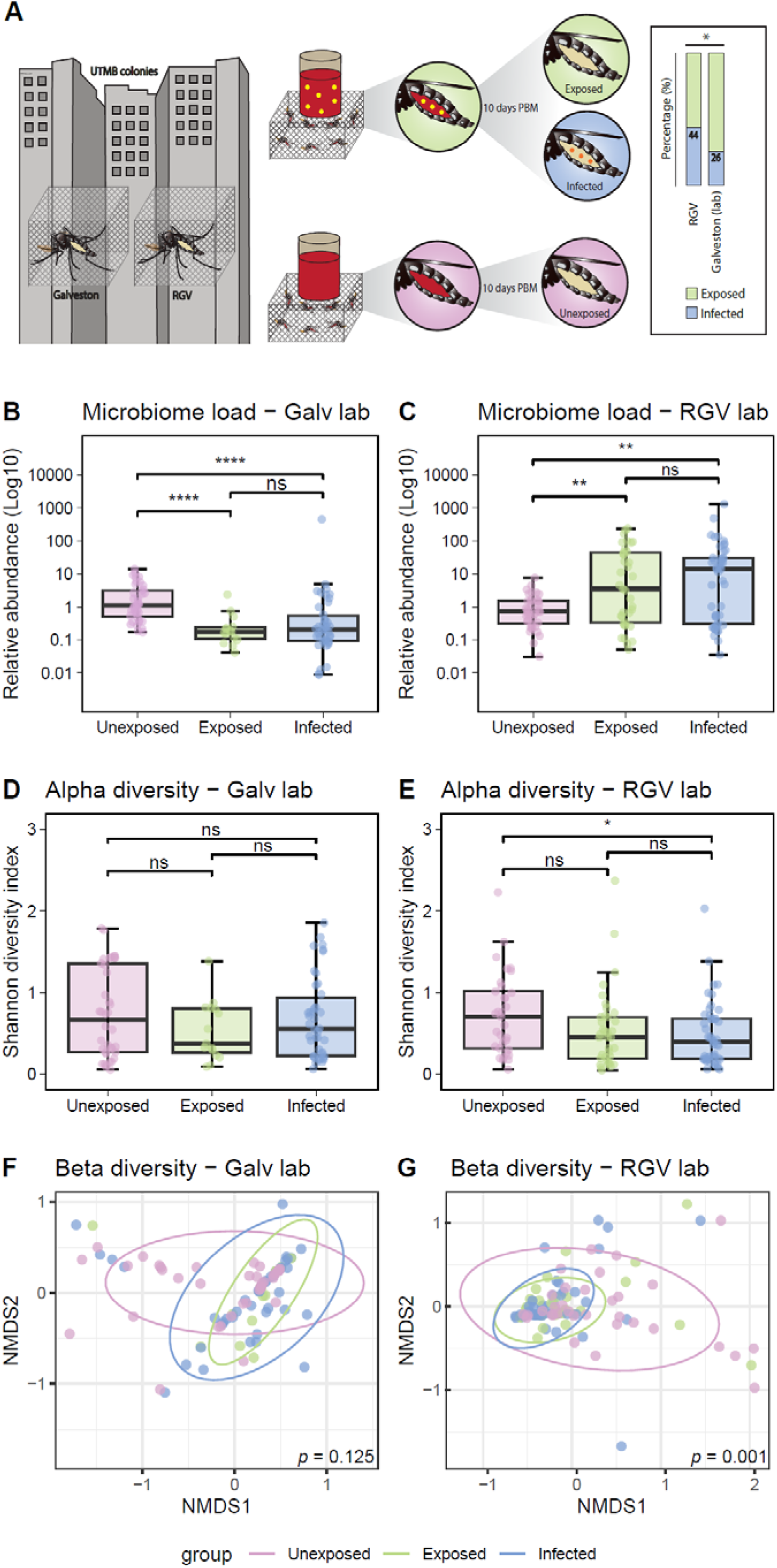
Viral infection of lab-reared mosquitoes and impact on the microbiome. Two *Ae. aegypti* lines reared in the insectaries of UTMB, Galveston (N=97) and Rio Grande Valley (RGV) (N=125), were offered a bloodmeal (red) spiked with ZIKV (yellow). Additionally, laboratory-reared mosquitoes were offered an uninfected bloodmeal (unexposed, pink). Ten days post bloodmeal (PBM) infection was assessed and mosquitoes were classified in exposed (ZIKV was not detected) (green) or infected (ZIKV was detected) (blue). Infection rate was assessed (right) and statistical difference is shown as * (Chi-square, p<0.05) (**A**). Relative abundance of bacterial 16S rRNA was measured in Galveston (**B**) and RGV (**C**) mosquitoes. Alpha diversity (Shannon diversity index) of the microbiome was assessed in Galveston (**D**) and RGV (**E**) mosquitoes. Statistical differences are shown as **** (*p*<0.0001), ** (*p*<0.01), * (*p*<0.05) and ns (non-significant) (Wilcoxon Rank Test). Beta diversity of the microbiome was assessed in Galveston (**F**) and RGV (**G**) mosquitoes. *p* values show results of PERMANOVA analysis of Bray-Curtis dissimilarity. Subsequent pairwise testing of beta diversity indicated in the RGV group, there were statistically significant differences between both unexposed vs. exposed and unexposed vs. infected (both *p*<0.003).

To assess whether ZIKV affected the microbiomes of these two distinct laboratory-reared mosquito lines in a similar fashion, we compared density, diversity, and composition of the microbiome among the three groups (unexposed, exposed, and infected) for each host line. In the Galveston-lab line, ZIKV exposure and infection led to a reduction in bacterial density compared to unexposed (Wilcoxon Rank Test, *p*<0.0001) (**Figure 2B**). Conversely, in the RGV-lab line, ZIKV exposure and infection resulted in an increase in bacterial density (Wilcoxon Rank Test, *p*<0.01) (**Figure 2C**). In the Galveston-lab line, neither ZIKV exposure nor infection caused significant differences in alpha or beta diversity (**Figure 2D, F**). However, ZIKV infection led to a significant reduction in Shannon’s diversity of the RGV lines microbiome (Wilcoxon Rank Test, *p*<0.05) (**Figure 2E**), while both exposure and infection significantly altered beta diversity compared to unexposed (PERMANOVA, *p*<0.01) (**Figure 2G**).

To evaluate whether the native microbiome was different between the two mosquito lines, we examined the diversity of the unaltered (ZIKV-unexposed) microbiome. While no significant difference was observed in alpha diversity between the lines (**Figure 3A**), beta diversity displayed a significant difference (PERMANOVA, *p*=0.006) (**Figure 3B**). These findings suggested that the differential impact of ZIKV on the RGV-lab and Galveston-lab lines may be attributed, at least partially, to the distinct composition of their microbiomes prior to infection. To elucidate whether ZIKV exposure and infection similarly affect microbiome composition in the two distinct lab lines, we characterised the microbiomes of unexposed, exposed and infected individuals in individuals from each line. Irrespective of ZIKV infection status, both host lines were dominated by *Acetobacteraceae* (**Figure 3C,D**) but members of the *Enterobacteriaceae* family were notable in the Galveston-lab line.

**Figure 3.**
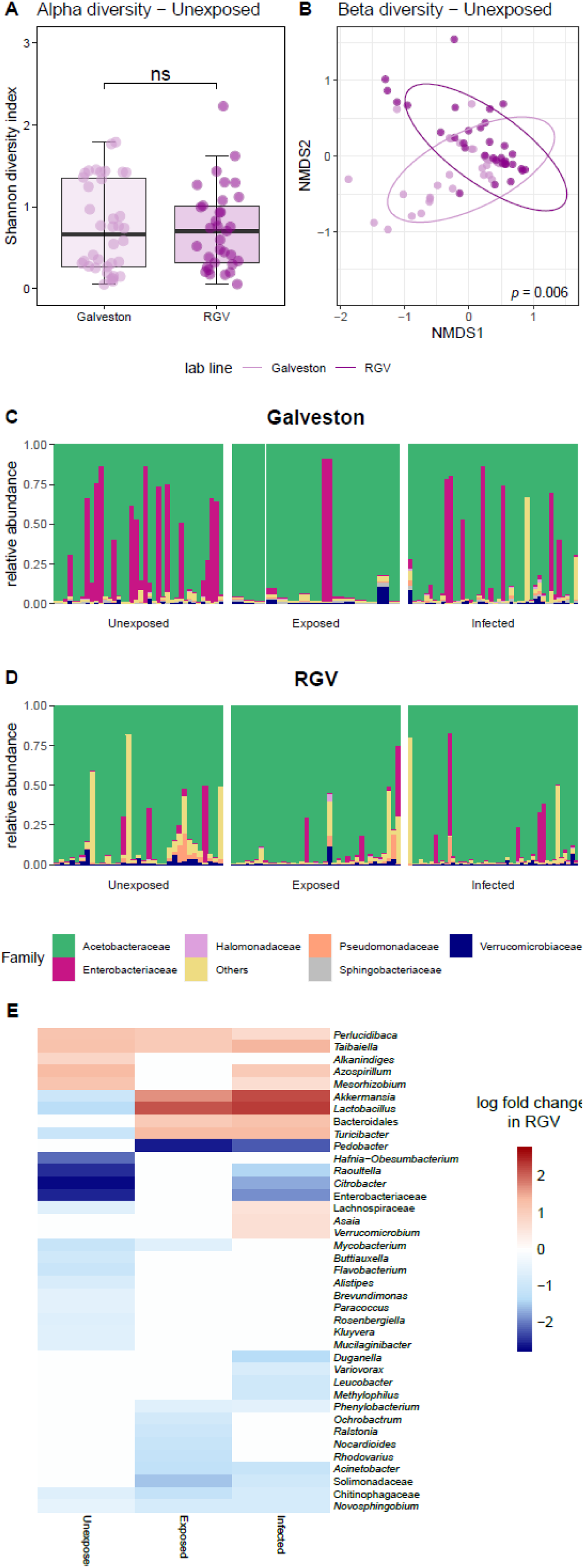
Comparison of microbiome diversity between *Ae. aegypti* laboratory lines. Alpha diversity (**A**) and beta diversity (**B**) were assessed in unexposed RGV and Galveston mosquitoes. Statistical differences are shown as ns (non-significant) (Wilcoxon Rank Test). *p* values show results of PERMANOVA analysis of Bray-Curtis dissimilarity. Relative abundance of bacterial families was explored in Galveston (**C**) and RGV (**D**) mosquitoes either unexposed, ZIKV exposed or ZIKV infected. The heatmap shows the ANCOM-BC results (adjusted *p*-value<0.05) of enriched taxa (red) or depleted taxa (blue) in RGV mosquitoes in comparison with Galveston mosquitoes within the unexposed, ZIKV-infected and ZIKV-exposed groups (**E**).

The two mosquito lines, which were derived from different regions, had distinct microbiome compositions, potentially leading to certain microbial taxa responding differently to ZIKV exposure and infection. To identify whether particular taxa show opposing trends between lines, we examined differential abundance in the microbiome composition between the Galveston-lab and RGV-lab lines, considering each condition. A total of 39 taxa exhibited significant differential abundance between the two lines when comparing each condition separately (**Figure 3E**). *Turicibacter*, *Akkermansia* and *Lactobacillus* showed the most pronounced changes. These bacteria had higher relative abundances in Galveston-lab mosquitoes in the unexposed cohort but this shifted in the infected and exposed groups with increases in the RGV-lab line. Conversely, both ZIKV exposure and infection resulted in a relative decrease of *Pedobacter* and *Acinetobacter* in RGV-lab mosquitoes compared to Galveston-lab mosquitoes. Taken together, these findings demonstrate the specific microbial taxa in distinct mosquito lines respond differently to ZIKV exposure and infection.

### Bacterial taxa correlate with ZIKV infection in *Ae. aegypti*

Next, we examined whether variation in the microbiome correlated to viral infection in the mosquito. We therefore examined the differential abundance of the microbiome, comparing the infection status (exposed and infected) in both the RGV-lab and Galveston-lab lines. We saw no differentially abundant bacteria in the RGV-lab line, while three bacteria were different in the Galveston-lab line; a *Rhizobium* and *Perlucidaca* were more prevalent in infected mosquitoes while *Ochrobactrum* was enriched in exposed mosquitoes (**Figure 4A**). To determine how the presence of the virus in the mosquito midgut shaped the microbiome, we also compared unexposed mosquitoes to both exposed and infected. Here we saw more profound effects with several taxa altered. In Galveston-lab mosquitoes, the majority of differentially abundant bacteria were more enriched in the unexposed group, and only *Ochrobactrum* and *Elizabethkingia* were enriched in the exposed group (**Figure 4B**). Four bacteria (*Tanticharoenia*, *Leucobacter, Enterobacter, Elizabethkingia)* were enriched in the infected group (**Figure 4C**). Conversely, the majority of taxa that showed significant changes in the RGV-lab line were enriched in the exposed or infected group compared to the unexposed control (**Figure 4D,E)**, further highlighting the distinction between these two lab lines.

**Figure 4.**
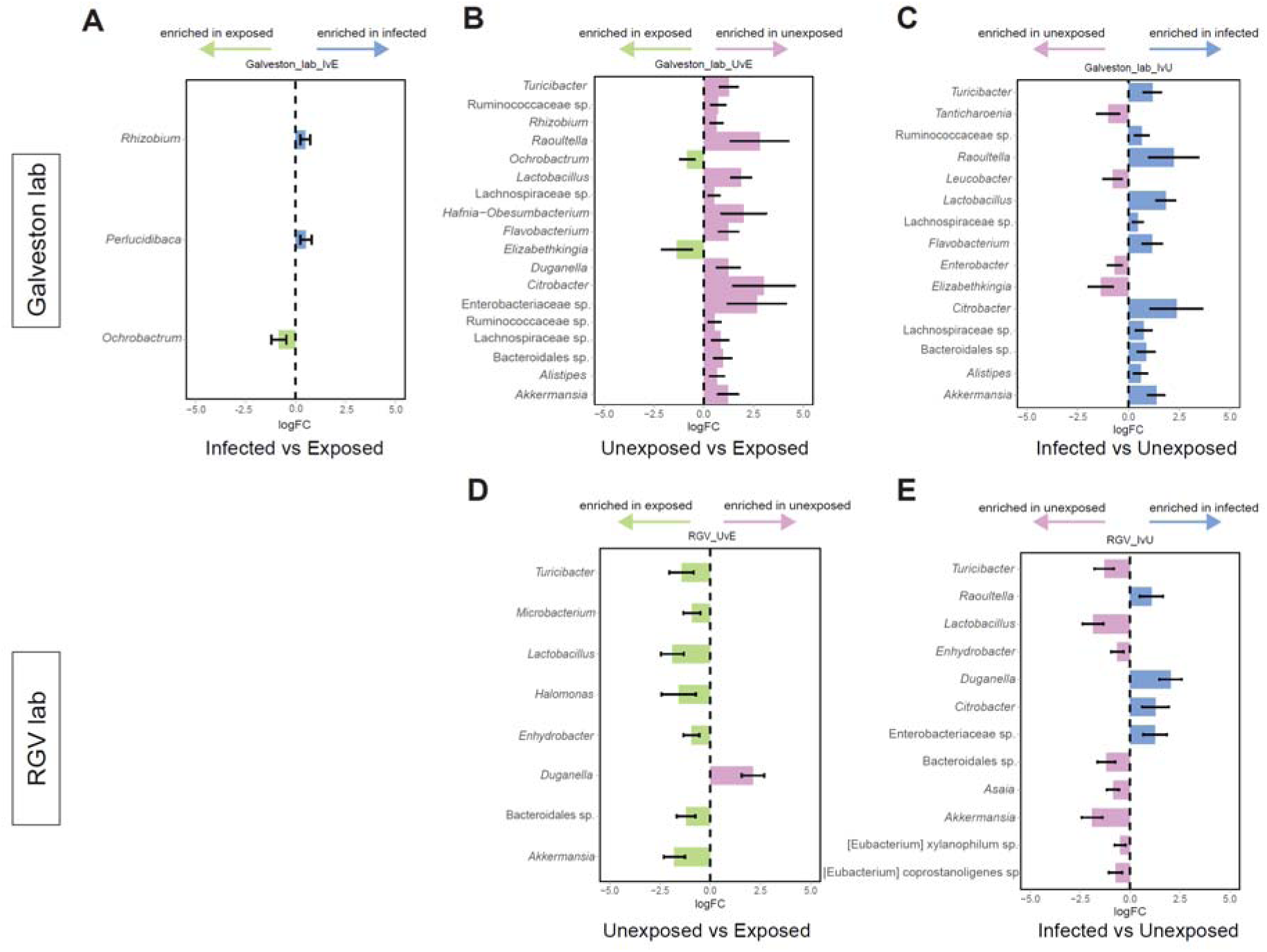
Differential abundance of microbes based on infection status. ANCOM (adjusted *p*-value <0.05) was used to identify taxa that were enriched in exposed (green) or infected (blue) Galveston mosquitoes (**A**). No differentially abundant taxa were identified in RGV mosquitoes. Differentially abundant taxa comparing unexposed to exposed (**B,D)** and unexposed to infected (**C,E**) in Galveston (**B,C**) and RGV (**D,E**) mosquitoes. Colours indicate taxa enriched in unexposed (pink), exposed (green) and infected (blue) mosquitoes.

### Microbiome-ZIKV interactions in field-collected mosquitoes

In order to ascertain whether our insights from laboratory findings would be representative of observations from field conditions, we examined if different mosquitoes collected from the field influenced progression of ZIKV infection. Host seeking *Ae. aegypti* mosquitoes were caught in three regions in Texas and immediately offered a blood meal spiked with ZIKV. After 10 days, virus infection status, microbiome composition and load were determined. Infection status was evaluated as done previously, whereby mosquitoes were categorised as exposed, if the virus did not progress, or infected if virus infection in the midgut could be determined. The prevalence of infection was comparable across sites, with infection rates recorded at 57%, 50% and 42% in mosquitoes collected in Austin, Galveston, and Brownsville, respectively (**Figure 5A**).

**Figure 5.**
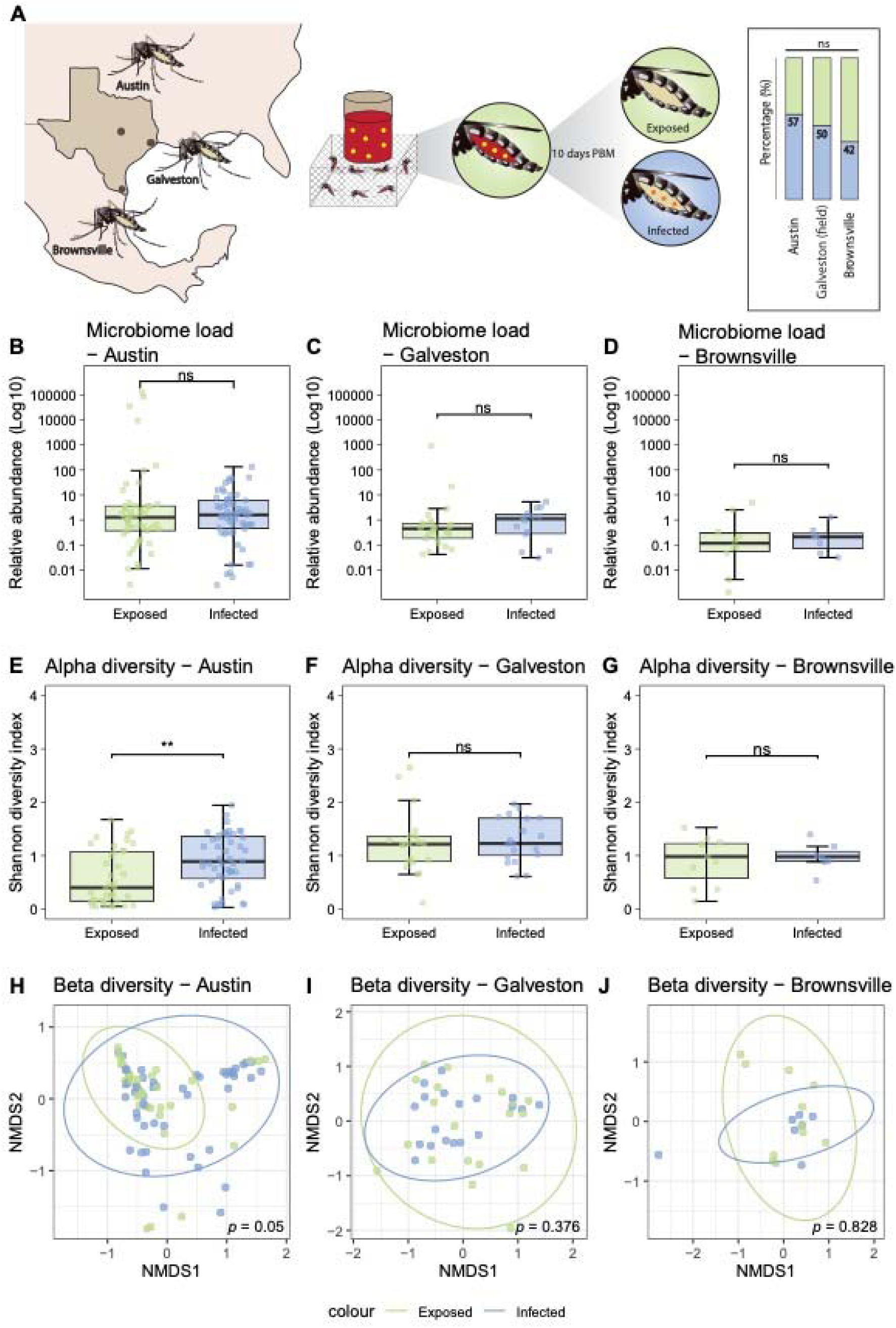
ZIKV infection of field-collected *Ae. aegypti* mosquitoes and impact of virus on the microbiome load and diversity. Field collected *Ae. aegypti* mosquitoes were collected from three locations in Texas; Austin (N=113), Galveston (N=40) and Brownsville (N=19), and offered a ZIKV infected blood meal. infection was assessed and mosquitoes were classified in exposed (ZIKV was not detected, green) or infected (ZIKV was detected, blue). Infection rate was assessed (right) and statistical difference is shown as * (Chi-square, *p*<0.05) (**A**). Relative abundance of bacterial 16S rRNA in Austin (**B**), Galveston (**C**) and Brownsville (**D**) mosquitoes. Alpha diversity (Shannon diversity index) of the microbiome in Austin (**E**), Galveston (**F**) and Brownsville (**G**) mosquitoes. Statistical differences are shown as ** (*p*<0.01) and ns (non-significant) (Wilcoxon rank test). Beta diversity of the microbiome in Austin (**H**), Galveston (**I**) and Brownsville (**J**) mosquitoes. Pairwise PERMANOVA was used for statistical analysis of the Bray-Curtis dissimilarity distance of microbiomes (bottom right of panel).

We first confirmed that the microbiome of field collected mosquitoes differed compared to their lab-counterparts by comparing Galveston-field to Galveston-lab mosquitoes from within the exposed or infected groups. Both the alpha and beta diversity was significantly different when comparing lab to field mosquitoes (**Figure S2)**. To further explore microbiome dynamics associated with ZIKV infection in field-collected mosquitoes, we examined the relative abundance of bacterial taxa in ZIKV-exposed and ZIKV-infected mosquitoes. Across all locations, no taxa were significantly differentially abundant when comparing infected and exposed groups. *Acetobacteraceae* represented the major microbiome component in Austin-field mosquitoes, while *Pseudomonadaceae* were more prevalent in Galveston-field mosquitoes (**Figure S3**).

To examine the impact of the microbiome on ZIKV infection in mosquitoes from three geographically distant locations, we conducted a comparative analysis of the microbiome between exposed and infected mosquitoes from each field site. We observed no differences in the bacterial load following viral infection in mosquitoes from any location (**Figure 5B-D**). However, when examining the diversity of the microbiome in exposed and infected mosquitoes from each location, significant differences in alpha (Wilcoxon Rank Test, *p*<0.01) and beta (PERMANOVA, *p*=0.04) diversity uniquely observed in mosquitoes collected from Austin (**Figure 5E-J**).

## Discussion

The microbiome of mosquitoes is highly variable and shaped by factors such as the environment, host, and microbial interactions [37, 38]. As such, mosquitoes of the same species collected in different geographical settings often harbour diverse microbiomes. Similarly, colonisation of mosquitoes alters their microbiome which is often less diverse compared to their field counterparts, while mosquitoes reared in distinct insectaries can exhibit considerable variation in their microbiome [4, 39]. It is therefore imperative to understand how microbiome variation influences vector competence and how universal these effects are between distinct mosquito lines. Here we show that ZIKV infection modulates the microbiome of mosquitoes in a host-line dependant manner. Importantly, we demonstrate this in both lab-reared and field-collected mosquitoes that have distinct microbiomes of differing complexity.

While a range of diverse arboviruses have been shown to alter the mosquito microbiome [6, 40–43], the effect on different mosquito lines had not yet been examined. We showed that viral exposure or infection of two lab colonies resulted in profoundly different microbial responses. An infectious blood meal reduced the total bacterial load of Galveston-lab mosquitoes yet increased load in the RGV-lab line. Similarly, we saw differences between the two lab lines in the alpha and beta diversity when comparing the unexposed to infected groups. In corroboration of our results for the RGV-lab group, Chikungunya virus (CHIKV) infection reduced alpha diversity in *Ae. aegpyti*; however, in contrast, another study has shown that both ZIKV and La Crosse virus (LACV) infection increased bacterial richness in *Ae. aegpyti*, *Ae. japonicus* and *Ae. triseriatus* [41, 42]. Importantly, we also found variable effects of viral infection and exposure on the microbiome in field collected samples. Infection altered both alpha and beta diversity of mosquito microbiomes collected in Austin but not those collected from Brownsville or Galveston. Our sampling was conducting in three regions in Texas, however more granular sampling may be required to examine within region differences in a mosquito population. To delve further into the difference seen in the lab-reared lines we examined bacterial taxa that differed between the viral exposed and infected groups which could accounts for the observed shifts in the microbiome. In the Galveston-lab line, bacteria including *Pedobacter*, *Enterobacter* and *Citrobacter* were significantly enriched in infected individuals, while *Lactobacillus*, *Akkermansia*, and *Turicibacter* were enriched in exposed and infected RGV-lab mosquitoes. Both *Enterobacter* and *Citrobacter* have been shown to increase in abundance after a CHIKV infection in *Aedes albopictus* mosquitoes [40, 44].

We were also interested in correlating microbes that were differentially abundant in infected compared to exposed individuals as these were potential microbes that could facilitate or interfere with infection respectively. Again, we saw distinct differences between the lines, with *Rhizobium* and *Perlucidibace* more prevalent in the infected while *Ochrobactrum* was more abundant in the exposed individuals in the Galveston-lab line, but no differentially abundant bacteria were found in the RGV-lab line. Little is known about these species in mosquitoes although *Ochrobactrum* has been associated with insecticide resistant mosquitoes [45]. In contrast, we saw no differentially abundant bacteria between exposed and infected groups in field collected mosquitoes. This could be related to these mosquitoes harbouring a more diverse microbiome or that life histories and age of field collected mosquitoes were unknown but likely less uniform compared to the lab-reared mosquitoes. Alternatively, it could be due to changes in the microbiome post viral exposure. In our experiments we assessed both ZIKV infection and the microbiome at 10 days post exposure to an infectious blood meal. However, the microbiome is dynamic and changes over the course of the mosquito’s life, and these changes may mask initial differences that influenced virus progression at the time of blood feeding [46]. Supporting this is the finding microbiome differences were less pronounced in ZIKV-infected mosquitoes at 21 compared to seven dpi, suggesting that microbiomes reverted toward the non-infectious state over time, potentially as the immune response returns to baseline or due to prolonged sugar feeding [41].

We also compared differentially abundant bacteria in unexposed mosquitoes to exposed and infected mosquitoes within a line. The bidirectionality of the system complicates understanding these interactions, as the presence of the microbe could affect pathogen progression or alternatively the presence of microbe may be indicative of their ability to persist within the pathogen-infected host compared to other members of the microbiome. Differences in bacterial abundance in the exposed group, whereby host immune pathways are triggered compared to the infected group, may be useful in differentiating between these scenarios. The Galveston-lab and RGV-lab mosquitoes were distinct regarding these differences, with the majority of bacterial taxa more abundant in the unexposed Galveston-lab group, whereas the reverse was the case for the RGV-lab line. *Akkermansia*, *Bacteroidales*, and *Turicibacter* had contrasting infection patterns. When looking at specific bacterial taxa that are more well known for in their interactions with mosquitoes, saw the *Elizabethkingia* was enriched in the Galveston-lab line in exposed and infected groups. *Elizabethkingia* has previously been shown to have ZIKV blocking potential and the identification of its presence here in exposed and infected mosquitoes provides credence to the comparative design to identify bacteria with anti-pathogen effects [47]. *Asaia* was enriched in the infected in the RGV-lab line. It’s dominance of the microbiome and known ability to influence pathogens makes it a candidate to further examine its influence on vector competence to ZIKV [48, 49]. *Tanticharoenia*, which belongs to the same family as *Asaia*, displayed a similar pattern to *Asaia* with greater abundance in the ZIKV infection Galveston-lab line.

While *Akkermansia* and *Turicibacter* are less well-known members of the mosquito microbiome, they have been observed in descriptive studies [43, 50, 51]. These bacteria are more recognized for their colonisation of mammalian guts and higher abundances of both these taxa were seen in the guts of *Plasmodium*-infected compared to uninfected mice, suggesting these bacteria are modulated by infection in general across diverse hosts [52–54]. While the mechanism(s) are unclear, it is known that *Turicibacter* is modulated by serotonin in vertebrates. In mosquitoes, ZIKV infection can alter serotonin levels of the neurotransmitter, Serotonin, so this could be a potentially under-explored mechanism by which infection alters the microbiome [55]. Further work is required to determine if distinct mosquito lines have differential serotonin responses to infection which could lead to microbiome variation in response to pathogens.

It is well established that pathogen infection or microbiota colonization elicits an immune response in the mosquito and, in turn, these immune pathways interfere and control gut-associated bacteria and arboviruses, respectively [15, 16]. To that end, it has been postulated that insect immune pathways evolved alongside microbes and are used to maintain homeostasis of the gut microbiota, and these processes are particularly important for mosquitoes as they are immersed within these microbes in the larval environment [56]. As such, there are intricate tripartite interactions at play whereby both pathogens and microbiome abundance and composition are modulated by one another’s presence. Therefore, differences in immune profiles, microbiome compositions, and susceptibility of microbiota to host pathways could potentially explain the differential responses of the microbiomes of distinct mosquito lines to viral infection. Distinct global transcription profiles are observed in different host backgrounds in response to viral infection or microbial colonization [57–61]. As such, the variable response to infection in the host could mediate divergent microbial outcomes. Further comparative studies examining the variation in the transcriptional response to infection in a controlled system, investigating how host pathways influence microbial composition, would likely provide insights to the mechanisms mediating variability seen in our studies.

Here, we employed an approach to exploit the natural variation in the microbiome in mosquitoes and correlated this to viral infection outcomes. Furthermore, our design investigated host-microbe-pathogen interactions without the need for artificial perturbation of the microbiome, which can have adverse effect on the host. However, we do appreciate there are caveats to our design which should be considered when interpreting our results. For example, while field caught mosquitoes have more biological relevant microbiomes, they do impose other challenges such as the unknown variables regarding their genetics, age, life history, exposure to pathogens, and previous blood feeding status. Our infection process required these adult mosquitoes to be housed in containment facilities, and the influence on the microbiome when of maintaining adults on sucrose in a lab-environment is not fully appreciated. Procedures which transplant field microbiomes to mosquitoes in the lab [62–64] could be used in conjunction with approaches here to overcome some of these caveats. Despite these challenges, our approach did illuminate our understanding of mosquito-microbiome-pathogen interactions.

In conclusion we show that exposure to, or infection with, ZIKV in *Ae. aegypti* lines alters their microbiome in distinct fashions. These differences were observed in both lab-reared and field-collected mosquitoes. Different bacterial taxa were modulated between mosquito lines which may be due to bacterial alteration of viral infection or the susceptibility of bacterial taxa after virus infection, which is likely mediated by host pathways. Our results highlight how variation of the microbiomes of mosquitoes needs to be considered for interpretation of lab-based experiments and implementation of microbial-based strategies for vector-borne disease.

## Acknowledgements

EH and GLH were jointly supported by the BBSRC (BB/V011278/1, BB/V011278/2 to EH and GLH). Additionally, the BBSRC supported GLH (BB/T001240/1), a Royal Society Wolfson Fellowship (RSWF\R1\180013), NIH grants (R21AI124452 and R21AI129507), the UKRI (20197 and 85336), the EPSRC (V043911/1) and the NIHR (NIHR2000907). SCW was supported by NIH grant R24 AI120942. GLH is affiliated to the National Institute for Health Research Health Protection Research Unit (NIHR HPRU) in Emerging and Zoonotic Infections at University of Liverpool in partnership with Public Health England (PHE), in collaboration with Liverpool School of Tropical Medicine and the University of Oxford. The views expressed are those of the authors and not necessarily those of the NHS, the NIHR, the Department of Health or Public Health England. CCU was supported by a Medical Research Council Doctoral Training Scholarship (MR/N013514/1) to the Liverpool School of Tropical Medicine.

## Author contributions

Conceptualization – GLH, MAS; Data curation MAS, CCU, GG, KK; Formal analysis - MAS, CCU, GG, KK, LEB, EH; Funding acquisition GLH, SCW, EH; Investigation MAS, ALW; Methodology MAS, GG, KK, GLH; Project administration SCW, GLH; Resources SCW, GLH, ALW EH; Software GG, KK; Supervision SCW, EH, GLH; Validation CCU, LEB, EH, GLH; Visualization CCU, LEB; Writing – original draft - MAS, CCU; Writing – review & editing - LEB, EH, GLH, SCW.

## Availability of data and materials

The datasets generated, analysed, and supporting the conclusions of this article are available at **PRJNA1113645**, the detailed per-sample accession numbers are in Table S1. The R code used to analyse the data and produce all figures is publicly available at https://github.com/grant-hughes-lab/Zika-microbiome-interactions under zenodo id https://doi.org/10.5281/zenodo.14786744.

## References

1. Guégan, M., et al., The mosquito holobiont: fresh insight into mosquito-microbiota interactions. Microbiome, 2018. 6: p. 1–17.

2. Cansado-Utrilla, C., et al., The microbiome and mosquito vectorial capacity: rich potential for discovery and translation. Microbiome, 2021. 9(1): p. 111.

3. Accoti, A., et al., The influence of the larval microbiome on susceptibility to Zika virus is mosquito genotype-dependent. PLoS Pathogens, 2023. 19(10): p. e1011727.

4. Brettell, L.E., et al., Mosquitoes reared in nearby insectaries within an institution in close spatial proximity possess significantly divergent microbiomes. Environmental Microbiology, 2024. 27(1): e70027

5. Seabourn, P.S., et al., Aedes albopictus microbiome derives from environmental sources and partitions across distinct host tissues. Microbiologyopen, 2023. 12(3): p. e1364.

6. Ramirez, J.L., et al., Reciprocal tripartite interactions between the Aedes aegypti midgut microbiota, innate immune system and dengue virus influences vector competence. PLoS Neglected Tropical Diseases, 2012. 6(3): p. e1561.

7. Hughes, G.L., et al., Native microbiome impedes vertical transmission of Wolbachia in Anopheles mosquitoes. Proceedings of the National Academy of Sciences, 2014. 111(34): p. 12498–12503.

8. Hegde, S., et al., Interkingdom interactions shape the fungal microbiome of mosquitoes. Animal Microbiome, 2024. 6(1): p. 11.

9. Kozlova, E.V., et al., Microbial interactions in the mosquito gut determine Serratia colonization and blood-feeding propensity. The ISME journal, 2021. 15(1): p. 93–108.

10. Tesh, R.B., D.J. Gubler, and L. Rosen, Variation among goegraphic strains of Aedes albopictus in susceptibility to infection with chikungunya virus. The American journal of Tropical Medicine and Hygiene, 1976. 25(2): p. 326–335.

11. Bennett, K.E., et al., Variation in vector competence for dengue 2 virus among 24 collections of Aedes aegypti from Mexico and the United States. American Journal of Tropical Medicine and Hygiene, 2002. 67(1): p. 85–92.

12. Roundy, C.M., et al., Variation in Aedes aegypti mosquito competence for Zika virus transmission. Emerging Infectious Diseases, 2017. 23(4): p. 625.

13. Kilpatrick, A.M., et al., Spatial and temporal variation in vector competence of Culex pipiens and Cx. restuans mosquitoes for West Nile virus. The American journal of Tropical Medicine and Hygiene, 2010. 83(3): p. 607.

14. Gubler, D.J. and L. Rosen, Variation among geographic strains of Aedes albopictus in susceptibility to infection with dengue viruses. The American journal of tropical Medicine and Hygiene, 1976. 25(2): p. 318–325.

15. Gabrieli, P., et al., Mosquito trilogy: microbiota, immunity and pathogens, and their implications for the control of disease transmission. Frontiers in Microbiology, 2021. 12: p. 630438.

16. Cai, J.A. and G.K. Christophides, Immune interactions between mosquitoes and microbes during midgut colonization. Current Opinion in Insect Science, 2024: p. 101195.

17. Wu, P., et al., A gut commensal bacterium promotes mosquito permissiveness to arboviruses. Cell Host & Microbe, 2019. 25(1): p. 101–112. e5.

18. Saraiva, R.G., et al., Chromobacterium spp. mediate their anti-Plasmodium activity through secretion of the histone deacetylase inhibitor romidepsin. Scientific reports, 2018. 8(1): p. 6176.

19. Zhang, L., et al., A naturally isolated symbiotic bacterium suppresses flavivirus transmission by Aedes mosquitoes. Science, 2024. 384(6693): p. eadn9524.

20. Ballard, J. and R. Melvin, Tetracycline treatment influences mitochondrial metabolism and mtDNA density two generations after treatment in Drosophila. Insect Molecular Biology, 2007. 16(6): p. 799–802.

21. Chabanol, E., et al., Antibiotic treatment in Anopheles coluzzii affects carbon and nitrogen metabolism. Pathogens, 2020. 9(9): p. 679.

22. Guerbois, M., et al., *Outbreak of Zika virus infection, Chiapas State, Mexico*, *2015, and first confirmed transmission by Aedes aegypti mosquitoes in the Americas*. The Journal of Infectious Diseases, 2016. 214(9): p. 1349–1356.

23. Hegde, S., et al., Microbiome interaction networks and community structure from laboratory-reared and field-collected Aedes aegypti, Aedes albopictus, and Culex quinquefasciatus mosquito vectors. Frontiers in Microbiology, 2018. 9: p. 405381.

24. Weisburg, W.G., et al., 16S ribosomal DNA amplification for phylogenetic study. Journal of bacteriology, 1991. 173(2): p. 697–703.

25. Isoe, J., et al., Defects in coatomer protein I (COPI) transport cause blood feeding-induced mortality in Yellow Fever mosquitoes. Proceedings of the National Academy of Sciences, 2011. 108(24): p. E211–E217.

26. Livak, K.J. and T.D. Schmittgen, *Analysis of relative gene expression data using real-time quantitative PCR and the 2−* ΔΔ*CT method*. methods, 2001. 25(4): p. 402–408.

27. Kassambara, A., *ggpubr:’ggplot2’based publication ready plots*. R package version, 2018: p. 2.

28. R Core Team, R: A language and environment for statistical computing.. 2023.

29. Klindworth, A., et al., Evaluation of general 16S ribosomal RNA gene PCR primers for classical and next-generation sequencing-based diversity studies. Nucleic acids research, 2013. 41(1): p. e1–e1.

30. Quast, C., et al., The SILVA ribosomal RNA gene database project: improved data processing and web-based tools. Nucleic acids research, 2012. 41(D1): p. D590–D596.

31. McMurdie, P.J. and S. Holmes, *phyloseq: an R package for reproducible interactive analysis and graphics of microbiome census data*. PloS one, 2013. 8(4): p. e61217.

32. Oksanen J, S.G., Blanchet F, Kindt R, Legendre P, Minchin P, O’Hara R, Solymos P, Stevens M, Szoecs E, Wagner H, Barbour M, Bedward M, Bolker B, Borcard D, Carvalho G, Chirico M, De Caceres M, Durand S, Evangelista H, FitzJohn R, Friendly M,, vegan: Community Ecology Package. 2022.

33. Arbizu, P.M., pairwiseAdonis: Pairwise Multilevel Comparison using Adonis. 2017.

34. Wickham, H., ggplot2. Wiley interdisciplinary reviews: computational statistics, 2011. 3(2): p. 180–185.

35. Lin, H. and S.D. Peddada, Analysis of compositions of microbiomes with bias correction. Nature Communications, 2020. 11(1): p. 3514.

36. Kolde, R. and M.R. Kolde, Package ‘pheatmap’. R package, 2015. 1(7): p. 790.

37. Hixson, B., R. Chen, and N. Buchon, Innate immunity in Aedes mosquitoes: from pathogen resistance to shaping the microbiota. Philosophical Transactions of the Royal Society B, 2024. 379(1901): p. 20230063.

38. Strand, M.R., Composition and functional roles of the gut microbiota in mosquitoes. Current opinion in insect science, 2018. 28: p. 59–65.

39. Accoti, A., et al., Variable microbiomes between mosquito lines are maintained across different environments. PLoS Neglected Tropical Diseases, 2023. 17(9): p. e0011306.

40. Zouache, K., et al., Chikungunya virus impacts the diversity of symbiotic bacteria in mosquito vector. Molecular Ecology, 2012. 21(9): p. 2297–2309.

41. Shi, C., et al., Bidirectional Interactions between Arboviruses and the Bacterial and Viral Microbiota in Aedes aegypti and Culex quinquefasciatus. MBio, 2022. 13(5): p. e01021–22.

42. Muturi, E.J., et al., Midgut fungal and bacterial microbiota of Aedes triseriatus and Aedes japonicus shift in response to La Crosse virus infection. Molecular Ecology, 2016. 25(16): p. 4075–4090.

43. Arévalo-Cortés, A., et al., Association of Midgut bacteria and their metabolic pathways with Zika infection and insecticide resistance in Colombian Aedes aegypti populations. Viruses, 2022. 14(10): p. 2197.

44. Siriyasatien, P., et al., Comparative analysis of midgut bacterial communities in Chikungunya virus-infected and non-infected Aedes aegypti Thai laboratory strain mosquitoes. Scientific Reports, 2024. 14(1): p. 10814.

45. Pelloquin, B., et al., Overabundance of Asaia and Serratia bacteria is associated with deltamethrin insecticide susceptibility in Anopheles coluzzii from Agboville, Côte d’Ivoire. Microbiology Spectrum, 2021. 9(2): p. e00157–21.

46. Wang, Y., et al., Dynamic gut microbiome across life history of the malaria mosquito Anopheles gambiae in Kenya. PloS one, 2011. 6(9): p. e24767.

47. Onyango, M.G., et al., Zika virus and temperature modulate Elizabethkingia anophelis in Aedes albopictus. Parasites & Vectors, 2021. 14: p. 1–15.

48. Tatsinkou Maffo, C.G., et al., Contrasting patterns of Asaia association with Plasmodium falciparum between field-collected Anopheles gambiae and Anopheles coluzzii from Cameroon. Microbiology Spectrum, 2024. 12(12): p. e00567–24.

49. Bassene, H., et al., 16S metagenomic comparison of Plasmodium falciparum–infected and noninfected Anopheles gambiae and Anopheles funestus microbiota from Senegal. The American Journal of Tropical Medicine and Hygiene, 2018. 99(6): p. 1489.

50. Rodpai, R., et al., Microbiome composition and microbial community structure in mosquito vectors Aedes aegypti and Aedes albopictus in Northeastern Thailand, a dengue-endemic area. Insects, 2023. 14(2): p. 184.

51. Sharma, P., et al., Salivary glands harbor more diverse microbial communities than gut in Anopheles culicifacies. Parasites & Vectors, 2014. 7: p. 1–7.

52. Belzer, C. and W.M. De Vos, Microbes inside—from diversity to function: the case of Akkermansia. The ISME journal, 2012. 6(8): p. 1449–1458.

53. Knowler, S.A., et al., Altered gastrointestinal tract structure and microbiome following cerebral malaria infection. Parasitology Research, 2023. 122(3): p. 789–799.

54. Guan, W., et al., Observation of intestinal flora diversity with the parasites infection process in a nonlethal malaria model of BALB/c mice induced by Plasmodium yoelii 17XNL strain. Decoding Infection and Transmission, 2023. 1: p. 100004.

55. Onyango, M.G., et al., Zika virus infection results in biochemical changes associated with RNA editing, inflammatory and antiviral responses in Aedes albopictus. Frontiers in Microbiology, 2020. 11: p. 559035.

56. Hanson, M.A., When the microbiome shapes the host: immune evolution implications for infectious disease. Philosophical Transactions of the Royal Society B, 2024. 379(1901): p. 20230061.

57. Etebari, K., et al., Global transcriptome analysis of Aedes aegypti mosquitoes in response to Zika virus infection. MSphere, 2017. 2(6): p. 10.1128/msphere.00456-17.

58. Hyde, J., et al., Limited influence of the microbiome on the transcriptional profile of female Aedes aegypti mosquitoes. Scientific reports, 2020. 10(1): p. 10880.

59. Jia, N., et al., Transcriptome Analysis of Response to Zika Virus Infection in Two Aedes albopictus Strains with Different Vector Competence. International Journal of Molecular Sciences, 2023. 24(5): p. 4257.

60. Vogel, K.J., et al., Transcriptome sequencing reveals large-scale changes in axenic Aedes aegypti larvae. PLoS Neglected Tropical Diseases, 2017. 11(1): p. e0005273.

61. Wang, S., et al., A cell atlas of the adult female Aedes aegypti midgut revealed by single-cell RNA sequencing. Scientific Data, 2024. 11(1): p. 587.

62. Coon, K.L., et al., Interspecies microbiome transplantation recapitulates microbial acquisition in mosquitoes. Microbiome 10, 58 (2022).

63. Zhao SY, et al., A cryopreservation method to recover laboratory- and field-derived bacterial communities from mosquito larval habitats. PLoS Neglected Tropical Diseases, 2023. 5;17(4):e0011234.

64. LaReau JC, et al., Introducing an environmental microbiome to axenic *Aedes aegypti* mosquitoes documents bacterial responses to a blood meal. Appl Environ Microbiol, 2023. 89:e00959–23.

