## Supplementary files for "Mosquito host background influences microbiome-ZIKV interactions in field and laboratory-reared *Aedes aegypti*"

Cansado-Utrilla et al. Supplementary files.

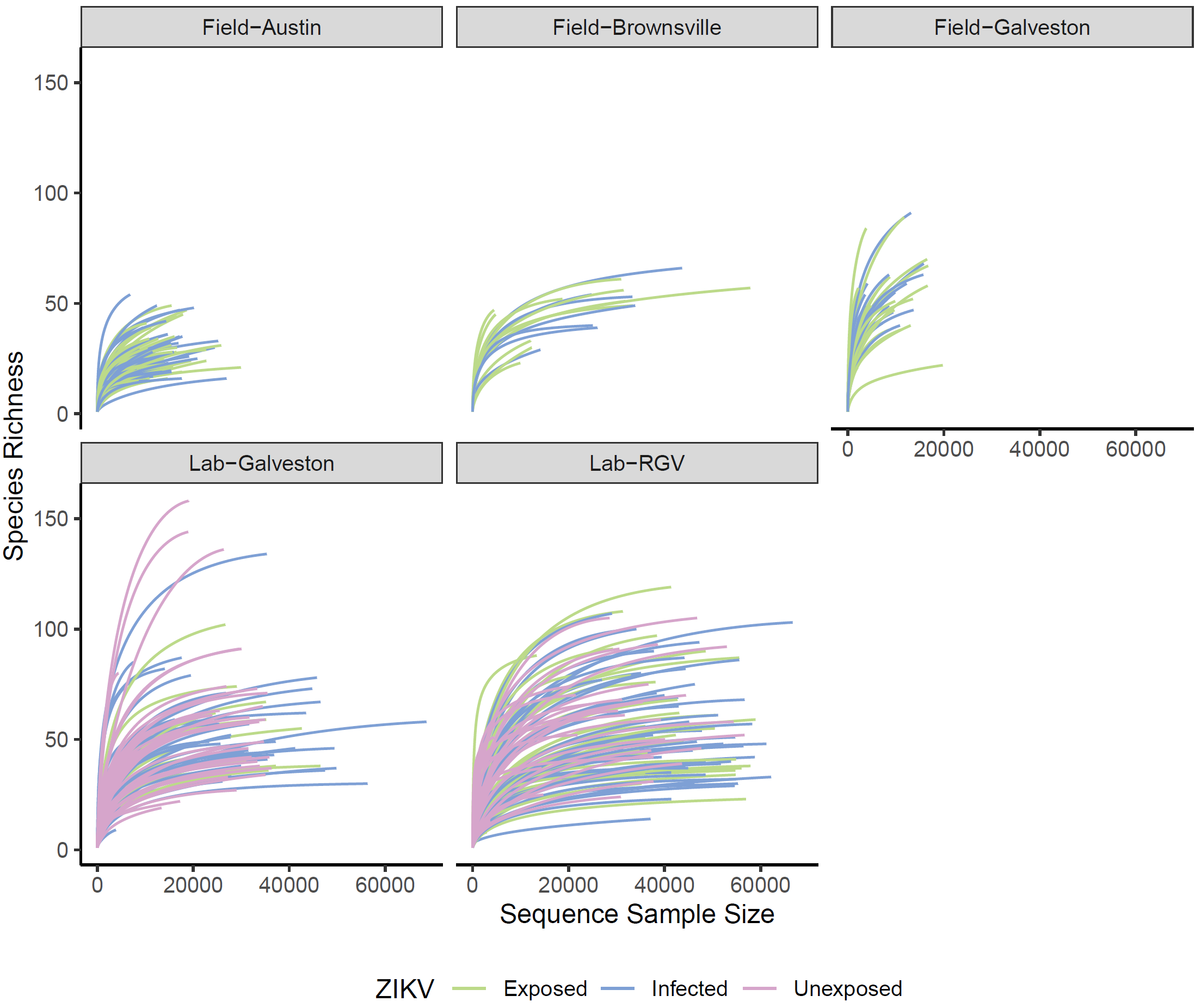

**Supplementary Figure 1. Rarefactions curves of all 16S samples facetted by sample type and coloured according to ZIKV infection status.** Samples with fewer than 2000 reads in total were removed prior to plotting.

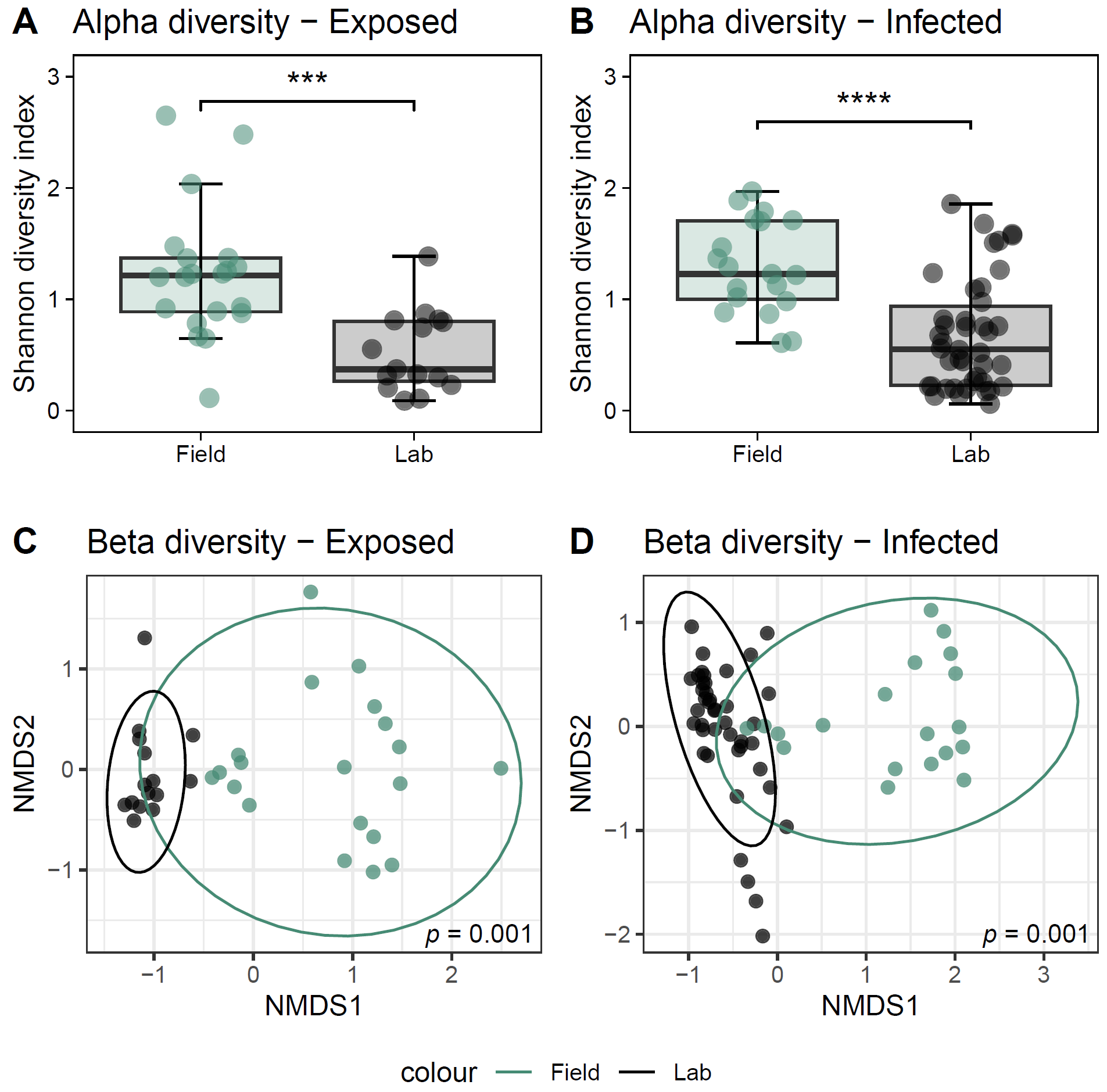

**Supplementary Figure 2. Comparison of microbiome diversity and composition between the laboratory-reared line originally derived from Galveston and mosquitoes freshly collected from field traps in Galveston.** ZIKV-exposed (A, C) and ZIKV-infected (B, D) mosquitoes were analysed separately. Statistical differences in alpha diversity are shown as **** (p<0.0001) and *** (p<0.001) (Wilcoxon Rank Test) (A, B). Pairwise PERMANOVA was used for statistical analysis of the Bray-Curtis dissimilarity distance of microbiomes (C, D). Beta diversity was significantly different between laboratory and field mosquitoes in the exposed and infected groups (Permanova, P=0.001).

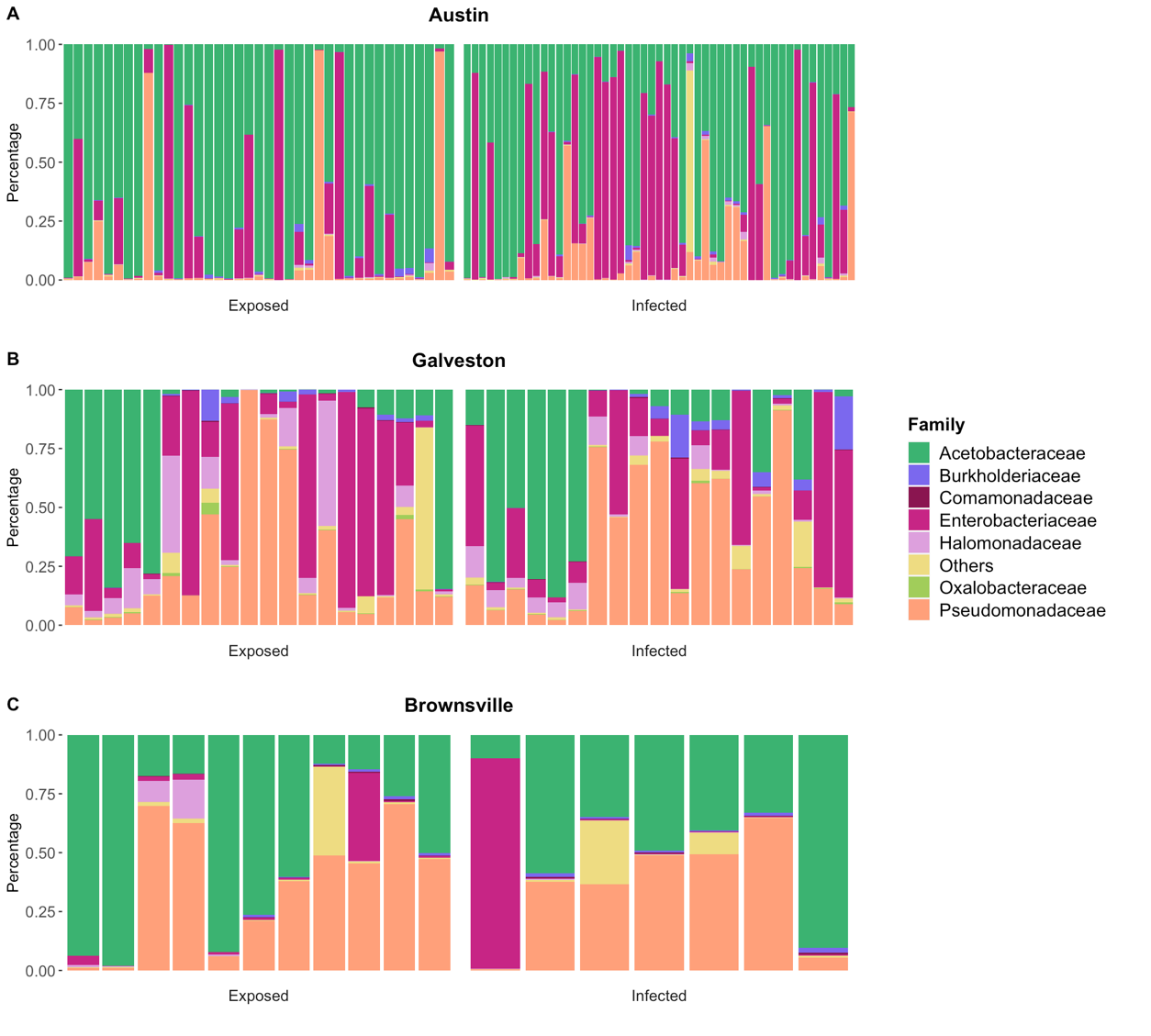

**Supplementary Figure 3. Impact of ZIKV on the relative abundance of microbial taxa in field-collected *Ae. aegypti* mosquitoes.** Relative abundance of bacterial families was explored in Austin (A), Galveston (B) and Brownsville (C) mosquitoes either ZIKV exposed or ZIKV infected.
