## Supplementary table for "Mosquito host background influences microbiome-ZIKV interactions in field and laboratory-reared *Aedes aegypti*"

Table S1. Metadata of the 16S rRNA sequencing samples.

| Sample ID | Reads | Type | ZIKV | Location | Sequencing Batch |
| --- | --- | --- | --- | --- | --- |
| Austin1-4_S4 | Austin1-4_S4_L001_R1_001 (paired).R1.fastq.gz  Austin1-4_S4_L001_R1_001 (paired).R2.fastq.gz | Field | Infected | Austin | Batch_1 |
| Austin1-5_S5 | Austin1-5_S5_L001_R1_001 (paired).R1.fastq.gz  Austin1-5_S5_L001_R1_001 (paired).R2.fastq.gz | Field | Infected | Austin | Batch_1 |
| Austin1-6_S6 | Austin1-6_S6_L001_R1_001 (paired).R1.fastq.gz  Austin1-6_S6_L001_R1_001 (paired).R2.fastq.gz | Field | Infected | Austin | Batch_1 |
| Austin1-7_S7 | Austin1-7_S7_L001_R1_001 (paired).R1.fastq.gz  Austin1-7_S7_L001_R1_001 (paired).R2.fastq.gz | Field | Infected | Austin | Batch_1 |
| Austin1-8_S8 | Austin1-8_S8_L001_R1_001 (paired).R1.fastq.gz  Austin1-8_S8_L001_R1_001 (paired).R2.fastq.gz | Field | Infected | Austin | Batch_1 |
| Austin1-9_S9 | Austin1-9_S9_L001_R1_001 (paired).R1.fastq.gz  Austin1-9_S9_L001_R1_001 (paired).R2.fastq.gz | Field | Infected | Austin | Batch_1 |
| Austin1-10_S10 | Austin1-10_S10_L001_R1_001 (paired).R1.fastq.gz  Austin1-10_S10_L001_R1_001 (paired).R2.fastq.gz | Field | Infected | Austin | Batch_1 |
| Austin1-11_S11 | Austin1-11_S11_L001_R1_001 (paired).R1.fastq.gz  Austin1-11_S11_L001_R1_001 (paired).R2.fastq.gz | Field | Exposed | Austin | Batch_1 |
| Austin1-12_S12 | Austin1-12_S12_L001_R1_001 (paired).R1.fastq.gz  Austin1-12_S12_L001_R1_001 (paired).R2.fastq.gz | Field | Infected | Austin | Batch_1 |
| Austin1-13_S13 | Austin1-13_S13_L001_R1_001 (paired).R1.fastq.gz  Austin1-13_S13_L001_R1_001 (paired).R2.fastq.gz | Field | Infected | Austin | Batch_1 |
| Austin1-14_S14 | Austin1-14_S14_L001_R1_001 (paired).R1.fastq.gz  Austin1-14_S14_L001_R1_001 (paired).R2.fastq.gz | Field | Exposed | Austin | Batch_1 |
| Austin1-15_S15 | Austin1-15_S15_L001_R1_001 (paired).R1.fastq.gz  Austin1-15_S15_L001_R1_001 (paired).R2.fastq.gz | Field | Infected | Austin | Batch_1 |
| Austin1-16_S16 | Austin1-16_S16_L001_R1_001 (paired).R1.fastq.gz  Austin1-16_S16_L001_R1_001 (paired).R2.fastq.gz | Field | Exposed | Austin | Batch_1 |
| Austin1-17_S17 | Austin1-17_S17_L001_R1_001 (paired).R1.fastq.gz  Austin1-17_S17_L001_R1_001 (paired).R2.fastq.gz | Field | Infected | Austin | Batch_1 |
| Austin1-18_S18 | Austin1-18_S18_L001_R1_001 (paired).R1.fastq.gz  Austin1-18_S18_L001_R1_001 (paired).R2.fastq.gz | Field | Exposed | Austin | Batch_1 |
| Austin1-19_S19 | Austin1-19_S19_L001_R1_001 (paired).R1.fastq.gz  Austin1-19_S19_L001_R1_001 (paired).R2.fastq.gz | Field | Exposed | Austin | Batch_1 |
| Austin2-1_S20 | Austin2-1_S20_L001_R1_001 (paired).R1.fastq.gz  Austin2-1_S20_L001_R1_001 (paired).R2.fastq.gz | Field | Infected | Austin | Batch_1 |
| Austin2-2_S21 | Austin2-2_S21_L001_R1_001 (paired).R1.fastq.gz  Austin2-2_S21_L001_R1_001 (paired).R2.fastq.gz | Field | Infected | Austin | Batch_1 |
| Austin2-3_S22 | Austin2-3_S22_L001_R1_001 (paired).R1.fastq.gz  Austin2-3_S22_L001_R1_001 (paired).R2.fastq.gz | Field | Exposed | Austin | Batch_1 |
| Austin2-4_S23 | Austin2-4_S23_L001_R1_001 (paired).R1.fastq.gz  Austin2-4_S23_L001_R1_001 (paired).R2.fastq.gz | Field | Infected | Austin | Batch_1 |
| Austin2-5_S24 | Austin2-5_S24_L001_R1_001 (paired).R1.fastq.gz  Austin2-5_S24_L001_R1_001 (paired).R2.fastq.gz | Field | Infected | Austin | Batch_1 |
| Austin2-6_S25 | Austin2-6_S25_L001_R1_001 (paired).R1.fastq.gz  Austin2-6_S25_L001_R1_001 (paired).R2.fastq.gz | Field | Infected | Austin | Batch_1 |
| Austin2-7_S26 | Austin2-7_S26_L001_R1_001 (paired).R1.fastq.gz  Austin2-7_S26_L001_R1_001 (paired).R2.fastq.gz | Field | Infected | Austin | Batch_1 |
| Austin2-8_S27 | Austin2-8_S27_L001_R1_001 (paired).R1.fastq.gz  Austin2-8_S27_L001_R1_001 (paired).R2.fastq.gz | Field | Exposed | Austin | Batch_1 |
| Austin2-9_S28 | Austin2-9_S28_L001_R1_001 (paired).R1.fastq.gz  Austin2-9_S28_L001_R1_001 (paired).R2.fastq.gz | Field | Infected | Austin | Batch_1 |
| Austin2-10_S29 | Austin2-10_S29_L001_R1_001 (paired).R1.fastq.gz  Austin2-10_S29_L001_R1_001 (paired).R2.fastq.gz | Field | Infected | Austin | Batch_1 |
| Austin2-11_S30 | Austin2-11_S30_L001_R1_001 (paired).R1.fastq.gz  Austin2-11_S30_L001_R1_001 (paired).R2.fastq.gz | Field | Exposed | Austin | Batch_1 |
| Austin2-12_S31 | Austin2-12_S31_L001_R1_001 (paired).R1.fastq.gz  Austin2-12_S31_L001_R1_001 (paired).R2.fastq.gz | Field | Infected | Austin | Batch_1 |
| Austin2-13_S32 | Austin2-13_S32_L001_R1_001 (paired).R1.fastq.gz  Austin2-13_S32_L001_R1_001 (paired).R2.fastq.gz | Field | Infected | Austin | Batch_1 |
| Austin2-14_S33 | Austin2-14_S33_L001_R1_001 (paired).R1.fastq.gz  Austin2-14_S33_L001_R1_001 (paired).R2.fastq.gz | Field | Exposed | Austin | Batch_1 |
| Austin2-15_S34 | Austin2-15_S34_L001_R1_001 (paired).R1.fastq.gz  Austin2-15_S34_L001_R1_001 (paired).R2.fastq.gz | Field | Infected | Austin | Batch_1 |
| Austin2-16_S35 | Austin2-16_S35_L001_R1_001 (paired).R1.fastq.gz  Austin2-16_S35_L001_R1_001 (paired).R2.fastq.gz | Field | Exposed | Austin | Batch_1 |
| Austin2-17_S36 | Austin2-17_S36_L001_R1_001 (paired).R1.fastq.gz  Austin2-17_S36_L001_R1_001 (paired).R2.fastq.gz | Field | Infected | Austin | Batch_1 |
| Austin2-18_S37 | Austin2-18_S37_L001_R1_001 (paired).R1.fastq.gz  Austin2-18_S37_L001_R1_001 (paired).R2.fastq.gz | Field | Exposed | Austin | Batch_1 |
| Austin2-19_S38 | Austin2-19_S38_L001_R1_001 (paired).R1.fastq.gz  Austin2-19_S38_L001_R1_001 (paired).R2.fastq.gz | Field | Exposed | Austin | Batch_1 |
| Austin2-20_S39 | Austin2-20_S39_L001_R1_001 (paired).R1.fastq.gz  Austin2-20_S39_L001_R1_001 (paired).R2.fastq.gz | Field | Infected | Austin | Batch_1 |
| Austin2-21_S40 | Austin2-21_S40_L001_R1_001 (paired).R1.fastq.gz  Austin2-21_S40_L001_R1_001 (paired).R2.fastq.gz | Field | Exposed | Austin | Batch_1 |
| Austin2-22_S41 | Austin2-22_S41_L001_R1_001 (paired).R1.fastq.gz  Austin2-22_S41_L001_R1_001 (paired).R2.fastq.gz | Field | Exposed | Austin | Batch_1 |
| Austin2-23_S42 | Austin2-23_S42_L001_R1_001 (paired).R1.fastq.gz  Austin2-23_S42_L001_R1_001 (paired).R2.fastq.gz | Field | Exposed | Austin | Batch_1 |
| Austin2A-1_S43 | Austin2A-1_S43_L001_R1_001 (paired).R1.fastq.gz  Austin2A-1_S43_L001_R1_001 (paired).R2.fastq.gz | Field | Exposed | Austin | Batch_1 |
| Austin2A-2_S44 | Austin2A-2_S44_L001_R1_001 (paired).R1.fastq.gz  Austin2A-2_S44_L001_R1_001 (paired).R2.fastq.gz | Field | Exposed | Austin | Batch_1 |
| Austin2A-3_S45 | Austin2A-3_S45_L001_R1_001 (paired).R1.fastq.gz  Austin2A-3_S45_L001_R1_001 (paired).R2.fastq.gz | Field | Infected | Austin | Batch_1 |
| Austin2A-4_S46 | Austin2A-4_S46_L001_R1_001 (paired).R1.fastq.gz  Austin2A-4_S46_L001_R1_001 (paired).R2.fastq.gz | Field | Exposed | Austin | Batch_1 |
| Austin2A-5_S47 | Austin2A-5_S47_L001_R1_001 (paired).R1.fastq.gz  Austin2A-5_S47_L001_R1_001 (paired).R2.fastq.gz | Field | Infected | Austin | Batch_1 |
| Austin2A-6_S48 | Austin2A-6_S48_L001_R1_001 (paired).R1.fastq.gz  Austin2A-6_S48_L001_R1_001 (paired).R2.fastq.gz | Field | Infected | Austin | Batch_1 |
| Austin2A-7_S49 | Austin2A-7_S49_L001_R1_001 (paired).R1.fastq.gz  Austin2A-7_S49_L001_R1_001 (paired).R2.fastq.gz | Field | Infected | Austin | Batch_1 |
| Austin2A-8_S50 | Austin2A-8_S50_L001_R1_001 (paired).R1.fastq.gz  Austin2A-8_S50_L001_R1_001 (paired).R2.fastq.gz | Field | Exposed | Austin | Batch_1 |
| Austin2A-9_S51 | Austin2A-9_S51_L001_R1_001 (paired).R1.fastq.gz  Austin2A-9_S51_L001_R1_001 (paired).R2.fastq.gz | Field | Exposed | Austin | Batch_1 |
| Austin2A-10_S52 | Austin2A-10_S52_L001_R1_001 (paired).R1.fastq.gz  Austin2A-10_S52_L001_R1_001 (paired).R2.fastq.gz | Field | Exposed | Austin | Batch_1 |
| Austin2A-11_S53 | Austin2A-11_S53_L001_R1_001 (paired).R1.fastq.gz  Austin2A-11_S53_L001_R1_001 (paired).R2.fastq.gz | Field | Exposed | Austin | Batch_1 |
| Austin2A-12_S54 | Austin2A-12_S54_L001_R1_001 (paired).R1.fastq.gz  Austin2A-12_S54_L001_R1_001 (paired).R2.fastq.gz | Field | Exposed | Austin | Batch_1 |
| Austin2A-13_S55 | Austin2A-13_S55_L001_R1_001 (paired).R1.fastq.gz  Austin2A-13_S55_L001_R1_001 (paired).R2.fastq.gz | Field | Exposed | Austin | Batch_1 |
| Austin2A-14_S56 | Austin2A-14_S56_L001_R1_001 (paired).R1.fastq.gz  Austin2A-14_S56_L001_R1_001 (paired).R2.fastq.gz | Field | Exposed | Austin | Batch_1 |
| Austin2A-15_S57 | Austin2A-15_S57_L001_R1_001 (paired).R1.fastq.gz  Austin2A-15_S57_L001_R1_001 (paired).R2.fastq.gz | Field | Exposed | Austin | Batch_1 |
| Austin2B-1_S58 | Austin2B-1_S58_L001_R1_001 (paired).R1.fastq.gz  Austin2B-1_S58_L001_R1_001 (paired).R2.fastq.gz | Field | Infected | Austin | Batch_1 |
| Austin2B-2_S59 | Austin2B-2_S59_L001_R1_001 (paired).R1.fastq.gz  Austin2B-2_S59_L001_R1_001 (paired).R2.fastq.gz | Field | Infected | Austin | Batch_1 |
| Austin2B-3_S60 | Austin2B-3_S60_L001_R1_001 (paired).R1.fastq.gz  Austin2B-3_S60_L001_R1_001 (paired).R2.fastq.gz | Field | Exposed | Austin | Batch_1 |
| Austin2B-4_S61 | Austin2B-4_S61_L001_R1_001 (paired).R1.fastq.gz  Austin2B-4_S61_L001_R1_001 (paired).R2.fastq.gz | Field | Infected | Austin | Batch_1 |
| Austin2B-5_S62 | Austin2B-5_S62_L001_R1_001 (paired).R1.fastq.gz  Austin2B-5_S62_L001_R1_001 (paired).R2.fastq.gz | Field | Infected | Austin | Batch_1 |
| Austin2B-6_S63 | Austin2B-6_S63_L001_R1_001 (paired).R1.fastq.gz  Austin2B-6_S63_L001_R1_001 (paired).R2.fastq.gz | Field | Exposed | Austin | Batch_1 |
| Austin2B-7_S64 | Austin2B-7_S64_L001_R1_001 (paired).R1.fastq.gz  Austin2B-7_S64_L001_R1_001 (paired).R2.fastq.gz | Field | Exposed | Austin | Batch_1 |
| Austin2B-8_S65 | Austin2B-8_S65_L001_R1_001 (paired).R1.fastq.gz  Austin2B-8_S65_L001_R1_001 (paired).R2.fastq.gz | Field | Infected | Austin | Batch_1 |
| Austin2B-9_S66 | Austin2B-9_S66_L001_R1_001 (paired).R1.fastq.gz  Austin2B-9_S66_L001_R1_001 (paired).R2.fastq.gz | Field | Infected | Austin | Batch_1 |
| Austin2B-10_S67 | Austin2B-10_S67_L001_R1_001 (paired).R1.fastq.gz  Austin2B-10_S67_L001_R1_001 (paired).R2.fastq.gz | Field | Infected | Austin | Batch_1 |
| Austin2B-11_S68 | Austin2B-11_S68_L001_R1_001 (paired).R1.fastq.gz  Austin2B-11_S68_L001_R1_001 (paired).R2.fastq.gz | Field | Infected | Austin | Batch_1 |
| Austin2B-12_S69 | Austin2B-12_S69_L001_R1_001 (paired).R1.fastq.gz  Austin2B-12_S69_L001_R1_001 (paired).R2.fastq.gz | Field | Infected | Austin | Batch_1 |
| Austin2B-13_S70 | Austin2B-13_S70_L001_R1_001 (paired).R1.fastq.gz  Austin2B-13_S70_L001_R1_001 (paired).R2.fastq.gz | Field | Infected | Austin | Batch_1 |
| Austin2B-14_S71 | Austin2B-14_S71_L001_R1_001 (paired).R1.fastq.gz  Austin2B-14_S71_L001_R1_001 (paired).R2.fastq.gz | Field | Infected | Austin | Batch_1 |
| Austin2B-15_S72 | Austin2B-15_S72_L001_R1_001 (paired).R1.fastq.gz  Austin2B-15_S72_L001_R1_001 (paired).R2.fastq.gz | Field | Exposed | Austin | Batch_1 |
| Austin3-1_S73 | Austin3-1_S73_L001_R1_001 (paired).R1.fastq.gz  Austin3-1_S73_L001_R1_001 (paired).R2.fastq.gz | Field | Infected | Austin | Batch_1 |
| Austin3-2_S74 | Austin3-2_S74_L001_R1_001 (paired).R1.fastq.gz  Austin3-2_S74_L001_R1_001 (paired).R2.fastq.gz | Field | Exposed | Austin | Batch_1 |
| Austin3-3_S75 | Austin3-3_S75_L001_R1_001 (paired).R1.fastq.gz  Austin3-3_S75_L001_R1_001 (paired).R2.fastq.gz | Field | Exposed | Austin | Batch_1 |
| Austin3-4_S76 | Austin3-4_S76_L001_R1_001 (paired).R1.fastq.gz  Austin3-4_S76_L001_R1_001 (paired).R2.fastq.gz | Field | Exposed | Austin | Batch_1 |
| Austin3-5_S77 | Austin3-5_S77_L001_R1_001 (paired).R1.fastq.gz  Austin3-5_S77_L001_R1_001 (paired).R2.fastq.gz | Field | Infected | Austin | Batch_1 |
| Austin3-6_S78 | Austin3-6_S78_L001_R1_001 (paired).R1.fastq.gz  Austin3-6_S78_L001_R1_001 (paired).R2.fastq.gz | Field | Exposed | Austin | Batch_1 |
| Austin3-7_S79 | Austin3-7_S79_L001_R1_001 (paired).R1.fastq.gz  Austin3-7_S79_L001_R1_001 (paired).R2.fastq.gz | Field | Exposed | Austin | Batch_1 |
| Austin3-8_S80 | Austin3-8_S80_L001_R1_001 (paired).R1.fastq.gz  Austin3-8_S80_L001_R1_001 (paired).R2.fastq.gz | Field | Infected | Austin | Batch_1 |
| Austin3-9_S81 | Austin3-9_S81_L001_R1_001 (paired).R1.fastq.gz  Austin3-9_S81_L001_R1_001 (paired).R2.fastq.gz | Field | Infected | Austin | Batch_1 |
| Austin3-10_S82 | Austin3-10_S82_L001_R1_001 (paired).R1.fastq.gz  Austin3-10_S82_L001_R1_001 (paired).R2.fastq.gz | Field | Exposed | Austin | Batch_1 |
| Austin3-11_S83 | Austin3-11_S83_L001_R1_001 (paired).R1.fastq.gz  Austin3-11_S83_L001_R1_001 (paired).R2.fastq.gz | Field | Infected | Austin | Batch_1 |
| Austin3-12_S84 | Austin3-12_S84_L001_R1_001 (paired).R1.fastq.gz  Austin3-12_S84_L001_R1_001 (paired).R2.fastq.gz | Field | Exposed | Austin | Batch_1 |
| Austin3-14_S86 | Austin3-14_S86_L001_R1_001 (paired).R1.fastq.gz  Austin3-14_S86_L001_R1_001 (paired).R2.fastq.gz | Field | Exposed | Austin | Batch_1 |
| Austin3-15_S87 | Austin3-15_S87_L001_R1_001 (paired).R1.fastq.gz  Austin3-15_S87_L001_R1_001 (paired).R2.fastq.gz | Field | Infected | Austin | Batch_1 |
| Austin3-16_S88 | Austin3-16_S88_L001_R1_001 (paired).R1.fastq.gz  Austin3-16_S88_L001_R1_001 (paired).R2.fastq.gz | Field | Infected | Austin | Batch_1 |
| Austin3-17_S89 | Austin3-17_S89_L001_R1_001 (paired).R1.fastq.gz  Austin3-17_S89_L001_R1_001 (paired).R2.fastq.gz | Field | Infected | Austin | Batch_1 |
| Austin3-18_S90 | Austin3-18_S90_L001_R1_001 (paired).R1.fastq.gz  Austin3-18_S90_L001_R1_001 (paired).R2.fastq.gz | Field | Exposed | Austin | Batch_1 |
| Austin3-19_S91 | Austin3-19_S91_L001_R1_001 (paired).R1.fastq.gz  Austin3-19_S91_L001_R1_001 (paired).R2.fastq.gz | Field | Infected | Austin | Batch_1 |
| Austin3-20_S92 | Austin3-20_S92_L001_R1_001 (paired).R1.fastq.gz  Austin3-20_S92_L001_R1_001 (paired).R2.fastq.gz | Field | Infected | Austin | Batch_1 |
| Austin3-21_S93 | Austin3-21_S93_L001_R1_001 (paired).R1.fastq.gz  Austin3-21_S93_L001_R1_001 (paired).R2.fastq.gz | Field | Infected | Austin | Batch_1 |
| Austin3-22_S94 | Austin3-22_S94_L001_R1_001 (paired).R1.fastq.gz  Austin3-22_S94_L001_R1_001 (paired).R2.fastq.gz | Field | Infected | Austin | Batch_1 |
| Austin3-23_S95 | Austin3-23_S95_L001_R1_001 (paired).R1.fastq.gz  Austin3-23_S95_L001_R1_001 (paired).R2.fastq.gz | Field | Exposed | Austin | Batch_1 |
| Austin3-24_S96 | Austin3-24_S96_L001_R1_001 (paired).R1.fastq.gz  Austin3-24_S96_L001_R1_001 (paired).R2.fastq.gz | Field | Exposed | Austin | Batch_1 |
| Austin2-1-1_S98 | Austin2-1-1_S98_L001_R1_001 (paired).R1.fastq.gz  Austin2-1-1_S98_L001_R1_001 (paired).R2.fastq.gz | Field | Infected | Austin | Batch_1 |
| Austin2-1-2_S99 | Austin2-1-2_S99_L001_R1_001 (paired).R1.fastq.gz  Austin2-1-2_S99_L001_R1_001 (paired).R2.fastq.gz | Field | Infected | Austin | Batch_1 |
| Austin2-1-3_S100 | Austin2-1-3_S100_L001_R1_001 (paired).R1.fastq.gz  Austin2-1-3_S100_L001_R1_001 (paired).R2.fastq.gz | Field | Exposed | Austin | Batch_1 |
| Austin2-1-4_S101 | Austin2-1-4_S101_L001_R1_001 (paired).R1.fastq.gz  Austin2-1-4_S101_L001_R1_001 (paired).R2.fastq.gz | Field | Exposed | Austin | Batch_1 |
| Austin2-1-5_S102 | Austin2-1-5_S102_L001_R1_001 (paired).R1.fastq.gz  Austin2-1-5_S102_L001_R1_001 (paired).R2.fastq.gz | Field | Infected | Austin | Batch_1 |
| Austin2-1-6_S103 | Austin2-1-6_S103_L001_R1_001 (paired).R1.fastq.gz  Austin2-1-6_S103_L001_R1_001 (paired).R2.fastq.gz | Field | Infected | Austin | Batch_1 |
| Austin2-1-7_S104 | Austin2-1-7_S104_L001_R1_001 (paired).R1.fastq.gz  Austin2-1-7_S104_L001_R1_001 (paired).R2.fastq.gz | Field | Infected | Austin | Batch_1 |
| Austin2-1-8_S105 | Austin2-1-8_S105_L001_R1_001 (paired).R1.fastq.gz  Austin2-1-8_S105_L001_R1_001 (paired).R2.fastq.gz | Field | Infected | Austin | Batch_1 |
| Austin2-2-1_S106 | Austin2-2-1_S106_L001_R1_001 (paired).R1.fastq.gz  Austin2-2-1_S106_L001_R1_001 (paired).R2.fastq.gz | Field | Infected | Austin | Batch_1 |
| Austin2-2-2_S107 | Austin2-2-2_S107_L001_R1_001 (paired).R1.fastq.gz  Austin2-2-2_S107_L001_R1_001 (paired).R2.fastq.gz | Field | Infected | Austin | Batch_1 |
| Austin2-2-3_S108 | Austin2-2-3_S108_L001_R1_001 (paired).R1.fastq.gz  Austin2-2-3_S108_L001_R1_001 (paired).R2.fastq.gz | Field | Exposed | Austin | Batch_1 |
| Austin2-2-4_S109 | Austin2-2-4_S109_L001_R1_001 (paired).R1.fastq.gz  Austin2-2-4_S109_L001_R1_001 (paired).R2.fastq.gz | Field | Infected | Austin | Batch_1 |
| Austin2-2-A_S110 | Austin2-2-A_S110_L001_R1_001 (paired).R1.fastq.gz  Austin2-2-A_S110_L001_R1_001 (paired).R2.fastq.gz | Field | Exposed | Austin | Batch_1 |
| Austin2-3-1_S111 | Austin2-3-1_S111_L001_R1_001 (paired).R1.fastq.gz  Austin2-3-1_S111_L001_R1_001 (paired).R2.fastq.gz | Field | Exposed | Austin | Batch_1 |
| Austin2-3-2_S112 | Austin2-3-2_S112_L001_R1_001 (paired).R1.fastq.gz  Austin2-3-2_S112_L001_R1_001 (paired).R2.fastq.gz | Field | Exposed | Austin | Batch_1 |
| Austin2-3-3_S113 | Austin2-3-3_S113_L001_R1_001 (paired).R1.fastq.gz  Austin2-3-3_S113_L001_R1_001 (paired).R2.fastq.gz | Field | Exposed | Austin | Batch_1 |
| GalvestonA-1_S114 | GalvestonA-1_S114_L001_R1_001 (paired).R1.fastq.gz  GalvestonA-1_S114_L001_R1_001 (paired).R2.fastq.gz | Field | Exposed | Galveston | Batch_1 |
| GalvestonA-2_S115 | GalvestonA-2_S115_L001_R1_001 (paired).R1.fastq.gz  GalvestonA-2_S115_L001_R1_001 (paired).R2.fastq.gz | Field | Exposed | Galveston | Batch_1 |
| GalvestonA-3_S116 | GalvestonA-3_S116_L001_R1_001 (paired).R1.fastq.gz  GalvestonA-3_S116_L001_R1_001 (paired).R2.fastq.gz | Field | Infected | Galveston | Batch_1 |
| GalvestonA-4_S117 | GalvestonA-4_S117_L001_R1_001 (paired).R1.fastq.gz  GalvestonA-4_S117_L001_R1_001 (paired).R2.fastq.gz | Field | Exposed | Galveston | Batch_1 |
| GalvestonA-5_S118 | GalvestonA-5_S118_L001_R1_001 (paired).R1.fastq.gz  GalvestonA-5_S118_L001_R1_001 (paired).R2.fastq.gz | Field | Exposed | Galveston | Batch_1 |
| GalvestonA-6_S119 | GalvestonA-6_S119_L001_R1_001 (paired).R1.fastq.gz  GalvestonA-6_S119_L001_R1_001 (paired).R2.fastq.gz | Field | Infected | Galveston | Batch_1 |
| GalvestonA-7_S120 | GalvestonA-7_S120_L001_R1_001 (paired).R1.fastq.gz  GalvestonA-7_S120_L001_R1_001 (paired).R2.fastq.gz | Field | Exposed | Galveston | Batch_1 |
| GalvestonA-8_S121 | GalvestonA-8_S121_L001_R1_001 (paired).R1.fastq.gz  GalvestonA-8_S121_L001_R1_001 (paired).R2.fastq.gz | Field | Infected | Galveston | Batch_1 |
| GalvestonA-9_S122 | GalvestonA-9_S122_L001_R1_001 (paired).R1.fastq.gz  GalvestonA-9_S122_L001_R1_001 (paired).R2.fastq.gz | Field | Infected | Galveston | Batch_1 |
| GalvestonA-10_S123 | GalvestonA-10_S123_L001_R1_001 (paired).R1.fastq.gz  GalvestonA-10_S123_L001_R1_001 (paired).R2.fastq.gz | Field | Infected | Galveston | Batch_1 |
| GalvestonA-11_S124 | GalvestonA-11_S124_L001_R1_001 (paired).R1.fastq.gz  GalvestonA-11_S124_L001_R1_001 (paired).R2.fastq.gz | Field | Infected | Galveston | Batch_1 |
| GalvestonB-1_S125 | GalvestonB-1_S125_L001_R1_001 (paired).R1.fastq.gz  GalvestonB-1_S125_L001_R1_001 (paired).R2.fastq.gz | Field | Exposed | Galveston | Batch_1 |
| GalvestonB-2_S126 | GalvestonB-2_S126_L001_R1_001 (paired).R1.fastq.gz  GalvestonB-2_S126_L001_R1_001 (paired).R2.fastq.gz | Field | Exposed | Galveston | Batch_1 |
| GalvestonB-3_S127 | GalvestonB-3_S127_L001_R1_001 (paired).R1.fastq.gz  GalvestonB-3_S127_L001_R1_001 (paired).R2.fastq.gz | Field | Exposed | Galveston | Batch_1 |
| GalvestonB-4_S128 | GalvestonB-4_S128_L001_R1_001 (paired).R1.fastq.gz  GalvestonB-4_S128_L001_R1_001 (paired).R2.fastq.gz | Field | Exposed | Galveston | Batch_1 |
| GalvestonB-5_S129 | GalvestonB-5_S129_L001_R1_001 (paired).R1.fastq.gz  GalvestonB-5_S129_L001_R1_001 (paired).R2.fastq.gz | Field | Exposed | Galveston | Batch_1 |
| GalvestonB-6_S130 | GalvestonB-6_S130_L001_R1_001 (paired).R1.fastq.gz  GalvestonB-6_S130_L001_R1_001 (paired).R2.fastq.gz | Field | Exposed | Galveston | Batch_1 |
| GalvestonB-7_S131 | GalvestonB-7_S131_L001_R1_001 (paired).R1.fastq.gz  GalvestonB-7_S131_L001_R1_001 (paired).R2.fastq.gz | Field | Exposed | Galveston | Batch_1 |
| GalvestonB-8_S132 | GalvestonB-8_S132_L001_R1_001 (paired).R1.fastq.gz  GalvestonB-8_S132_L001_R1_001 (paired).R2.fastq.gz | Field | Exposed | Galveston | Batch_1 |
| GalvestonB-9_S133 | GalvestonB-9_S133_L001_R1_001 (paired).R1.fastq.gz  GalvestonB-9_S133_L001_R1_001 (paired).R2.fastq.gz | Field | Infected | Galveston | Batch_1 |
| GalvestonB-10_S134 | GalvestonB-10_S134_L001_R1_001 (paired).R1.fastq.gz  GalvestonB-10_S134_L001_R1_001 (paired).R2.fastq.gz | Field | Infected | Galveston | Batch_1 |
| GalvestonB-11_S135 | GalvestonB-11_S135_L001_R1_001 (paired).R1.fastq.gz  GalvestonB-11_S135_L001_R1_001 (paired).R2.fastq.gz | Field | Exposed | Galveston | Batch_1 |
| GalvestonB-12_S136 | GalvestonB-12_S136_L001_R1_001 (paired).R1.fastq.gz  GalvestonB-12_S136_L001_R1_001 (paired).R2.fastq.gz | Field | Exposed | Galveston | Batch_1 |
| GalvestonB-13_S137 | GalvestonB-13_S137_L001_R1_001 (paired).R1.fastq.gz  GalvestonB-13_S137_L001_R1_001 (paired).R2.fastq.gz | Field | Infected | Galveston | Batch_1 |
| GalvestonB-14_S138 | GalvestonB-14_S138_L001_R1_001 (paired).R1.fastq.gz  GalvestonB-14_S138_L001_R1_001 (paired).R2.fastq.gz | Field | Exposed | Galveston | Batch_1 |
| Galveston2A-1_S139 | Galveston2A-1_S139_L001_R1_001 (paired).R1.fastq.gz  Galveston2A-1_S139_L001_R1_001 (paired).R2.fastq.gz | Field | Infected | Galveston | Batch_1 |
| Galveston2A-2_S140 | Galveston2A-2_S140_L001_R1_001 (paired).R1.fastq.gz  Galveston2A-2_S140_L001_R1_001 (paired).R2.fastq.gz | Field | Infected | Galveston | Batch_1 |
| Galveston2A-3_S141 | Galveston2A-3_S141_L001_R1_001 (paired).R1.fastq.gz  Galveston2A-3_S141_L001_R1_001 (paired).R2.fastq.gz | Field | Infected | Galveston | Batch_1 |
| Galveston2A-4_S142 | Galveston2A-4_S142_L001_R1_001 (paired).R1.fastq.gz  Galveston2A-4_S142_L001_R1_001 (paired).R2.fastq.gz | Field | Infected | Galveston | Batch_1 |
| Galveston2A-5_S143 | Galveston2A-5_S143_L001_R1_001 (paired).R1.fastq.gz  Galveston2A-5_S143_L001_R1_001 (paired).R2.fastq.gz | Field | Exposed | Galveston | Batch_1 |
| Galveston2A-6_S144 | Galveston2A-6_S144_L001_R1_001 (paired).R1.fastq.gz  Galveston2A-6_S144_L001_R1_001 (paired).R2.fastq.gz | Field | Infected | Galveston | Batch_1 |
| Galveston2A-7_S145 | Galveston2A-7_S145_L001_R1_001 (paired).R1.fastq.gz  Galveston2A-7_S145_L001_R1_001 (paired).R2.fastq.gz | Field | Exposed | Galveston | Batch_1 |
| Galveston2A-8_S146 | Galveston2A-8_S146_L001_R1_001 (paired).R1.fastq.gz  Galveston2A-8_S146_L001_R1_001 (paired).R2.fastq.gz | Field | Infected | Galveston | Batch_1 |
| Galveston2A-9_S147 | Galveston2A-9_S147_L001_R1_001 (paired).R1.fastq.gz  Galveston2A-9_S147_L001_R1_001 (paired).R2.fastq.gz | Field | Infected | Galveston | Batch_1 |
| Galveston2A-10_S148 | Galveston2A-10_S148_L001_R1_001 (paired).R1.fastq.gz  Galveston2A-10_S148_L001_R1_001 (paired).R2.fastq.gz | Field | Infected | Galveston | Batch_1 |
| Galveston2A-11_S149 | Galveston2A-11_S149_L001_R1_001 (paired).R1.fastq.gz  Galveston2A-11_S149_L001_R1_001 (paired).R2.fastq.gz | Field | Infected | Galveston | Batch_1 |
| Galveston2A-12_S150 | Galveston2A-12_S150_L001_R1_001 (paired).R1.fastq.gz  Galveston2A-12_S150_L001_R1_001 (paired).R2.fastq.gz | Field | Infected | Galveston | Batch_1 |
| Galveston2A-13_S151 | Galveston2A-13_S151_L001_R1_001 (paired).R1.fastq.gz  Galveston2A-13_S151_L001_R1_001 (paired).R2.fastq.gz | Field | Exposed | Galveston | Batch_1 |
| Galveston2A-14_S152 | Galveston2A-14_S152_L001_R1_001 (paired).R1.fastq.gz  Galveston2A-14_S152_L001_R1_001 (paired).R2.fastq.gz | Field | Infected | Galveston | Batch_1 |
| Galveston2B-1_S153 | Galveston2B-1_S153_L001_R1_001 (paired).R1.fastq.gz  Galveston2B-1_S153_L001_R1_001 (paired).R2.fastq.gz | Field | Exposed | Galveston | Batch_1 |
| Brownsville1_S154 | Brownsville1_S154_L001_R1_001 (paired).R1.fastq.gz  Brownsville1_S154_L001_R1_001 (paired).R2.fastq.gz | Field | Infected | Brownsville | Batch_1 |
| Brownsville2_S155 | Brownsville2_S155_L001_R1_001 (paired).R1.fastq.gz  Brownsville2_S155_L001_R1_001 (paired).R2.fastq.gz | Field | Exposed | Brownsville | Batch_1 |
| Brownsville3_S156 | Brownsville3_S156_L001_R1_001 (paired).R1.fastq.gz  Brownsville3_S156_L001_R1_001 (paired).R2.fastq.gz | Field | Exposed | Brownsville | Batch_1 |
| Brownsville4_S157 | Brownsville4_S157_L001_R1_001 (paired).R1.fastq.gz  Brownsville4_S157_L001_R1_001 (paired).R2.fastq.gz | Field | Exposed | Brownsville | Batch_1 |
| Brownsville5_S158 | Brownsville5_S158_L001_R1_001 (paired).R1.fastq.gz  Brownsville5_S158_L001_R1_001 (paired).R2.fastq.gz | Field | Infected | Brownsville | Batch_1 |
| Brownsville6_S159 | Brownsville6_S159_L001_R1_001 (paired).R1.fastq.gz  Brownsville6_S159_L001_R1_001 (paired).R2.fastq.gz | Field | Exposed | Brownsville | Batch_1 |
| Brownsville7_S160 | Brownsville7_S160_L001_R1_001 (paired).R1.fastq.gz  Brownsville7_S160_L001_R1_001 (paired).R2.fastq.gz | Field | Exposed | Brownsville | Batch_1 |
| Brownsville8_141_S141 | Brownsville8_141_S141_L001_R1_001 (paired).R1.fastq.gz  Brownsville8_141_S141_L001_R1_001 (paired).R2.fastq.gz | Field | Exposed | Brownsville | Batch_2 |
| Brownsville9_142_S142 | Brownsville9_142_S142_L001_R1_001 (paired).R1.fastq.gz  Brownsville9_142_S142_L001_R1_001 (paired).R2.fastq.gz | Field | Infected | Brownsville | Batch_2 |
| Brownsville10_143_S143 | Brownsville10_143_S143_L001_R1_001 (paired).R1.fastq.gz  Brownsville10_143_S143_L001_R1_001 (paired).R2.fastq.gz | Field | Exposed | Brownsville | Batch_2 |
| Brownsville11_144_S144 | Brownsville11_144_S144_L001_R1_001 (paired).R1.fastq.gz  Brownsville11_144_S144_L001_R1_001 (paired).R2.fastq.gz | Field | Infected | Brownsville | Batch_2 |
| Brownsville12_145_S145 | Brownsville12_145_S145_L001_R1_001 (paired).R1.fastq.gz  Brownsville12_145_S145_L001_R1_001 (paired).R2.fastq.gz | Field | Exposed | Brownsville | Batch_2 |
| Brownsville13_146_S146 | Brownsville13_146_S146_L001_R1_001 (paired).R1.fastq.gz  Brownsville13_146_S146_L001_R1_001 (paired).R2.fastq.gz | Field | Infected | Brownsville | Batch_2 |
| Brownsville14_147_S147 | Brownsville14_147_S147_L001_R1_001 (paired).R1.fastq.gz  Brownsville14_147_S147_L001_R1_001 (paired).R2.fastq.gz | Field | Infected | Brownsville | Batch_2 |
| Brownsville15_148_S148 | Brownsville15_148_S148_L001_R1_001 (paired).R1.fastq.gz  Brownsville15_148_S148_L001_R1_001 (paired).R2.fastq.gz | Field | Exposed | Brownsville | Batch_2 |
| Brownsville16_149_S149 | Brownsville16_149_S149_L001_R1_001 (paired).R1.fastq.gz  Brownsville16_149_S149_L001_R1_001 (paired).R2.fastq.gz | Field | Infected | Brownsville | Batch_2 |
| Brownsville17_150_S150 | Brownsville17_150_S150_L001_R1_001 (paired).R1.fastq.gz  Brownsville17_150_S150_L001_R1_001 (paired).R2.fastq.gz | Field | Exposed | Brownsville | Batch_2 |
| Brownsville18_151_S151 | Brownsville18_151_S151_L001_R1_001 (paired).R1.fastq.gz  Brownsville18_151_S151_L001_R1_001 (paired).R2.fastq.gz | Field | Exposed | Brownsville | Batch_2 |
| Brownsville19_152_S152 | Brownsville19_152_S152_L001_R1_001 (paired).R1.fastq.gz  Brownsville19_152_S152_L001_R1_001 (paired).R2.fastq.gz | Field | Infected | Brownsville | Batch_2 |
| Rio-Grande-Valley-Lab-Ae-aegypti-1_S1 | Rio-Grande-Valley-Lab-Ae-aegypti-1_S1_L001_R1_001 (paired).R1.fastq.gz  Rio-Grande-Valley-Lab-Ae-aegypti-1_S1_L001_R1_001 (paired).R2.fastq.gz | Lab | Exposed | RGV | Batch_1 |
| Rio-Grande-Valley-Lab-Ae-aegypti-2_S2 | Rio-Grande-Valley-Lab-Ae-aegypti-2_S2_L001_R1_001 (paired).R1.fastq.gz  Rio-Grande-Valley-Lab-Ae-aegypti-2_S2_L001_R1_001 (paired).R2.fastq.gz | Lab | Infected | RGV | Batch_1 |
| Rio-Grande-Valley-Lab-Ae-aegypti-3_S3 | Rio-Grande-Valley-Lab-Ae-aegypti-3_S3_L001_R1_001 (paired).R1.fastq.gz  Rio-Grande-Valley-Lab-Ae-aegypti-3_S3_L001_R1_001 (paired).R2.fastq.gz | Lab | Infected | RGV | Batch_1 |
| Rio-Grande-Valley-Lab-Ae-aegypti-4_S4 | Rio-Grande-Valley-Lab-Ae-aegypti-4_S4_L001_R1_001 (paired).R1.fastq.gz  Rio-Grande-Valley-Lab-Ae-aegypti-4_S4_L001_R1_001 (paired).R2.fastq.gz | Lab | Exposed | RGV | Batch_1 |
| Rio-Grande-Valley-Lab-Ae-aegypti-5_S5 | Rio-Grande-Valley-Lab-Ae-aegypti-5_S5_L001_R1_001 (paired).R1.fastq.gz  Rio-Grande-Valley-Lab-Ae-aegypti-5_S5_L001_R1_001 (paired).R2.fastq.gz | Lab | Infected | RGV | Batch_1 |
| Rio-Grande-Valley-Lab-Ae-aegypti-6_S6 | Rio-Grande-Valley-Lab-Ae-aegypti-6_S6_L001_R1_001 (paired).R1.fastq.gz  Rio-Grande-Valley-Lab-Ae-aegypti-6_S6_L001_R1_001 (paired).R2.fastq.gz | Lab | Infected | RGV | Batch_1 |
| Rio-Grande-Valley-Lab-Ae-aegypti-7_S7 | Rio-Grande-Valley-Lab-Ae-aegypti-7_S7_L001_R1_001 (paired).R1.fastq.gz  Rio-Grande-Valley-Lab-Ae-aegypti-7_S7_L001_R1_001 (paired).R2.fastq.gz | Lab | Infected | RGV | Batch_1 |
| Rio-Grande-Valley-Lab-Ae-aegypti-8_S8 | Rio-Grande-Valley-Lab-Ae-aegypti-8_S8_L001_R1_001 (paired).R1.fastq.gz  Rio-Grande-Valley-Lab-Ae-aegypti-8_S8_L001_R1_001 (paired).R2.fastq.gz | Lab | Infected | RGV | Batch_1 |
| Rio-Grande-Valley-Lab-Ae-aegypti-9_S9 | Rio-Grande-Valley-Lab-Ae-aegypti-9_S9_L001_R1_001 (paired).R1.fastq.gz  Rio-Grande-Valley-Lab-Ae-aegypti-9_S9_L001_R1_001 (paired).R2.fastq.gz | Lab | Infected | RGV | Batch_1 |
| Rio-Grande-Valley-Lab-Ae-aegypti-10_S10 | Rio-Grande-Valley-Lab-Ae-aegypti-10_S10_L001_R1_001 (paired).R1.fastq.gz  Rio-Grande-Valley-Lab-Ae-aegypti-10_S10_L001_R1_001 (paired).R2.fastq.gz | Lab | Infected | RGV | Batch_1 |
| Rio-Grande-Valley-Lab-Ae-aegypti-11_S11 | Rio-Grande-Valley-Lab-Ae-aegypti-11_S11_L001_R1_001 (paired).R1.fastq.gz  Rio-Grande-Valley-Lab-Ae-aegypti-11_S11_L001_R1_001 (paired).R2.fastq.gz | Lab | Infected | RGV | Batch_1 |
| Rio-Grande-Valley-Lab-Ae-aegypti-12_S12 | Rio-Grande-Valley-Lab-Ae-aegypti-12_S12_L001_R1_001 (paired).R1.fastq.gz  Rio-Grande-Valley-Lab-Ae-aegypti-12_S12_L001_R1_001 (paired).R2.fastq.gz | Lab | Infected | RGV | Batch_1 |
| Rio-Grande-Valley-Lab-Ae-aegypti-13_S13 | Rio-Grande-Valley-Lab-Ae-aegypti-13_S13_L001_R1_001 (paired).R1.fastq.gz  Rio-Grande-Valley-Lab-Ae-aegypti-13_S13_L001_R1_001 (paired).R2.fastq.gz | Lab | Exposed | RGV | Batch_1 |
| Rio-Grande-Valley-Lab-Ae-aegypti-14_S14 | Rio-Grande-Valley-Lab-Ae-aegypti-14_S14_L001_R1_001 (paired).R1.fastq.gz  Rio-Grande-Valley-Lab-Ae-aegypti-14_S14_L001_R1_001 (paired).R2.fastq.gz | Lab | Exposed | RGV | Batch_1 |
| Rio-Grande-Valley-Lab-Ae-aegypti-15_S15 | Rio-Grande-Valley-Lab-Ae-aegypti-15_S15_L001_R1_001 (paired).R1.fastq.gz  Rio-Grande-Valley-Lab-Ae-aegypti-15_S15_L001_R1_001 (paired).R2.fastq.gz | Lab | Infected | RGV | Batch_1 |
| Rio-Grande-Valley-Lab-Ae-aegypti-16_S16 | Rio-Grande-Valley-Lab-Ae-aegypti-16_S16_L001_R1_001 (paired).R1.fastq.gz  Rio-Grande-Valley-Lab-Ae-aegypti-16_S16_L001_R1_001 (paired).R2.fastq.gz | Lab | Exposed | RGV | Batch_1 |
| Rio-Grande-Valley-Lab-Ae-aegypti-17_S17 | Rio-Grande-Valley-Lab-Ae-aegypti-17_S17_L001_R1_001 (paired).R1.fastq.gz  Rio-Grande-Valley-Lab-Ae-aegypti-17_S17_L001_R1_001 (paired).R2.fastq.gz | Lab | Exposed | RGV | Batch_1 |
| Rio-Grande-Valley-Lab-Ae-aegypti-18_S18 | Rio-Grande-Valley-Lab-Ae-aegypti-18_S18_L001_R1_001 (paired).R1.fastq.gz  Rio-Grande-Valley-Lab-Ae-aegypti-18_S18_L001_R1_001 (paired).R2.fastq.gz | Lab | Infected | RGV | Batch_1 |
| Rio-Grande-Valley-Lab-Ae-aegypti-19_S19 | Rio-Grande-Valley-Lab-Ae-aegypti-19_S19_L001_R1_001 (paired).R1.fastq.gz  Rio-Grande-Valley-Lab-Ae-aegypti-19_S19_L001_R1_001 (paired).R2.fastq.gz | Lab | Infected | RGV | Batch_1 |
| Rio-Grande-Valley-Lab-Ae-aegypti-20_S20 | Rio-Grande-Valley-Lab-Ae-aegypti-20_S20_L001_R1_001 (paired).R1.fastq.gz  Rio-Grande-Valley-Lab-Ae-aegypti-20_S20_L001_R1_001 (paired).R2.fastq.gz | Lab | Exposed | RGV | Batch_1 |
| Rio-Grande-Valley-Lab-Ae-aegypti-21_S21 | Rio-Grande-Valley-Lab-Ae-aegypti-21_S21_L001_R1_001 (paired).R1.fastq.gz  Rio-Grande-Valley-Lab-Ae-aegypti-21_S21_L001_R1_001 (paired).R2.fastq.gz | Lab | Infected | RGV | Batch_1 |
| Rio-Grande-Valley-Lab-Ae-aegypti-22_S22 | Rio-Grande-Valley-Lab-Ae-aegypti-22_S22_L001_R1_001 (paired).R1.fastq.gz  Rio-Grande-Valley-Lab-Ae-aegypti-22_S22_L001_R1_001 (paired).R2.fastq.gz | Lab | Infected | RGV | Batch_1 |
| Rio-Grande-Valley-Lab-Ae-aegypti-23_S23 | Rio-Grande-Valley-Lab-Ae-aegypti-23_S23_L001_R1_001 (paired).R1.fastq.gz  Rio-Grande-Valley-Lab-Ae-aegypti-23_S23_L001_R1_001 (paired).R2.fastq.gz | Lab | Exposed | RGV | Batch_1 |
| Rio-Grande-Valley-Lab-Ae-aegypti-24_S24 | Rio-Grande-Valley-Lab-Ae-aegypti-24_S24_L001_R1_001 (paired).R1.fastq.gz  Rio-Grande-Valley-Lab-Ae-aegypti-24_S24_L001_R1_001 (paired).R2.fastq.gz | Lab | Exposed | RGV | Batch_1 |
| Rio-Grande-Valley-Lab-Ae-aegypti-25_S25 | Rio-Grande-Valley-Lab-Ae-aegypti-25_S25_L001_R1_001 (paired).R1.fastq.gz  Rio-Grande-Valley-Lab-Ae-aegypti-25_S25_L001_R1_001 (paired).R2.fastq.gz | Lab | Infected | RGV | Batch_1 |
| Rio-Grande-Valley-Lab-Ae-aegypti-26_S26 | Rio-Grande-Valley-Lab-Ae-aegypti-26_S26_L001_R1_001 (paired).R1.fastq.gz  Rio-Grande-Valley-Lab-Ae-aegypti-26_S26_L001_R1_001 (paired).R2.fastq.gz | Lab | Infected | RGV | Batch_1 |
| Rio-Grande-Valley-Lab-Ae-aegypti-27_S27 | Rio-Grande-Valley-Lab-Ae-aegypti-27_S27_L001_R1_001 (paired).R1.fastq.gz  Rio-Grande-Valley-Lab-Ae-aegypti-27_S27_L001_R1_001 (paired).R2.fastq.gz | Lab | Exposed | RGV | Batch_1 |
| Rio-Grande-Valley-Lab-Ae-aegypti-28_S28 | Rio-Grande-Valley-Lab-Ae-aegypti-28_S28_L001_R1_001 (paired).R1.fastq.gz  Rio-Grande-Valley-Lab-Ae-aegypti-28_S28_L001_R1_001 (paired).R2.fastq.gz | Lab | Exposed | RGV | Batch_1 |
| Rio-Grande-Valley-Lab-Ae-aegypti-29_S29 | Rio-Grande-Valley-Lab-Ae-aegypti-29_S29_L001_R1_001 (paired).R1.fastq.gz  Rio-Grande-Valley-Lab-Ae-aegypti-29_S29_L001_R1_001 (paired).R2.fastq.gz | Lab | Infected | RGV | Batch_1 |
| Rio-Grande-Valley-Lab-Ae-aegypti-30_S30 | Rio-Grande-Valley-Lab-Ae-aegypti-30_S30_L001_R1_001 (paired).R1.fastq.gz  Rio-Grande-Valley-Lab-Ae-aegypti-30_S30_L001_R1_001 (paired).R2.fastq.gz | Lab | Exposed | RGV | Batch_1 |
| Rio-Grande-Valley-Lab-Ae-aegypti-31_S31 | Rio-Grande-Valley-Lab-Ae-aegypti-31_S31_L001_R1_001 (paired).R1.fastq.gz  Rio-Grande-Valley-Lab-Ae-aegypti-31_S31_L001_R1_001 (paired).R2.fastq.gz | Lab | Infected | RGV | Batch_1 |
| Rio-Grande-Valley-Lab-Ae-aegypti-32_S32 | Rio-Grande-Valley-Lab-Ae-aegypti-32_S32_L001_R1_001 (paired).R1.fastq.gz  Rio-Grande-Valley-Lab-Ae-aegypti-32_S32_L001_R1_001 (paired).R2.fastq.gz | Lab | Infected | RGV | Batch_1 |
| Rio-Grande-Valley-Lab-Ae-aegypti-33_S33 | Rio-Grande-Valley-Lab-Ae-aegypti-33_S33_L001_R1_001 (paired).R1.fastq.gz  Rio-Grande-Valley-Lab-Ae-aegypti-33_S33_L001_R1_001 (paired).R2.fastq.gz | Lab | Infected | RGV | Batch_1 |
| Rio-Grande-Valley-Lab-Ae-aegypti-34_S34 | Rio-Grande-Valley-Lab-Ae-aegypti-34_S34_L001_R1_001 (paired).R1.fastq.gz  Rio-Grande-Valley-Lab-Ae-aegypti-34_S34_L001_R1_001 (paired).R2.fastq.gz | Lab | Exposed | RGV | Batch_1 |
| Rio-Grande-Valley-Lab-Ae-aegypti-35_S35 | Rio-Grande-Valley-Lab-Ae-aegypti-35_S35_L001_R1_001 (paired).R1.fastq.gz  Rio-Grande-Valley-Lab-Ae-aegypti-35_S35_L001_R1_001 (paired).R2.fastq.gz | Lab | Exposed | RGV | Batch_1 |
| Rio-Grande-Valley-Lab-Ae-aegypti-36_S36 | Rio-Grande-Valley-Lab-Ae-aegypti-36_S36_L001_R1_001 (paired).R1.fastq.gz  Rio-Grande-Valley-Lab-Ae-aegypti-36_S36_L001_R1_001 (paired).R2.fastq.gz | Lab | Exposed | RGV | Batch_1 |
| Rio-Grande-Valley-Lab-Ae-aegypti-37_S37 | Rio-Grande-Valley-Lab-Ae-aegypti-37_S37_L001_R1_001 (paired).R1.fastq.gz  Rio-Grande-Valley-Lab-Ae-aegypti-37_S37_L001_R1_001 (paired).R2.fastq.gz | Lab | Infected | RGV | Batch_1 |
| Rio-Grande-Valley-Lab-Ae-aegypti-38_S38 | Rio-Grande-Valley-Lab-Ae-aegypti-38_S38_L001_R1_001 (paired).R1.fastq.gz  Rio-Grande-Valley-Lab-Ae-aegypti-38_S38_L001_R1_001 (paired).R2.fastq.gz | Lab | Infected | RGV | Batch_1 |
| Rio-Grande-Valley-Lab-Ae-aegypti-39_S39 | Rio-Grande-Valley-Lab-Ae-aegypti-39_S39_L001_R1_001 (paired).R1.fastq.gz  Rio-Grande-Valley-Lab-Ae-aegypti-39_S39_L001_R1_001 (paired).R2.fastq.gz | Lab | Infected | RGV | Batch_1 |
| Rio-Grande-Valley-Lab-Ae-aegypti-40_S40 | Rio-Grande-Valley-Lab-Ae-aegypti-40_S40_L001_R1_001 (paired).R1.fastq.gz  Rio-Grande-Valley-Lab-Ae-aegypti-40_S40_L001_R1_001 (paired).R2.fastq.gz | Lab | Infected | RGV | Batch_1 |
| Rio-Grande-Valley-Lab-Ae-aegypti-41_S41 | Rio-Grande-Valley-Lab-Ae-aegypti-41_S41_L001_R1_001 (paired).R1.fastq.gz  Rio-Grande-Valley-Lab-Ae-aegypti-41_S41_L001_R1_001 (paired).R2.fastq.gz | Lab | Exposed | RGV | Batch_1 |
| Rio-Grande-Valley-Lab-Ae-aegypti-42_S42 | Rio-Grande-Valley-Lab-Ae-aegypti-42_S42_L001_R1_001 (paired).R1.fastq.gz  Rio-Grande-Valley-Lab-Ae-aegypti-42_S42_L001_R1_001 (paired).R2.fastq.gz | Lab | Exposed | RGV | Batch_1 |
| Rio-Grande-Valley-Lab-Ae-aegypti-43_S43 | Rio-Grande-Valley-Lab-Ae-aegypti-43_S43_L001_R1_001 (paired).R1.fastq.gz  Rio-Grande-Valley-Lab-Ae-aegypti-43_S43_L001_R1_001 (paired).R2.fastq.gz | Lab | Exposed | RGV | Batch_1 |
| Rio-Grande-Valley-Lab-Ae-aegypti-44_S44 | Rio-Grande-Valley-Lab-Ae-aegypti-44_S44_L001_R1_001 (paired).R1.fastq.gz  Rio-Grande-Valley-Lab-Ae-aegypti-44_S44_L001_R1_001 (paired).R2.fastq.gz | Lab | Infected | RGV | Batch_1 |
| Rio-Grande-Valley-Lab-Ae-aegypti-45_S45 | Rio-Grande-Valley-Lab-Ae-aegypti-45_S45_L001_R1_001 (paired).R1.fastq.gz  Rio-Grande-Valley-Lab-Ae-aegypti-45_S45_L001_R1_001 (paired).R2.fastq.gz | Lab | Infected | RGV | Batch_1 |
| Rio-Grande-Valley-Lab-Ae-aegypti-46_S46 | Rio-Grande-Valley-Lab-Ae-aegypti-46_S46_L001_R1_001 (paired).R1.fastq.gz  Rio-Grande-Valley-Lab-Ae-aegypti-46_S46_L001_R1_001 (paired).R2.fastq.gz | Lab | Infected | RGV | Batch_1 |
| Rio-Grande-Valley-Lab-Ae-aegypti-47_S47 | Rio-Grande-Valley-Lab-Ae-aegypti-47_S47_L001_R1_001 (paired).R1.fastq.gz  Rio-Grande-Valley-Lab-Ae-aegypti-47_S47_L001_R1_001 (paired).R2.fastq.gz | Lab | Exposed | RGV | Batch_1 |
| Rio-Grande-Valley-Lab-Ae-aegypti-48_S48 | Rio-Grande-Valley-Lab-Ae-aegypti-48_S48_L001_R1_001 (paired).R1.fastq.gz  Rio-Grande-Valley-Lab-Ae-aegypti-48_S48_L001_R1_001 (paired).R2.fastq.gz | Lab | Exposed | RGV | Batch_1 |
| Rio-Grande-Valley-Lab-Ae-aegypti-49_S49 | Rio-Grande-Valley-Lab-Ae-aegypti-49_S49_L001_R1_001 (paired).R1.fastq.gz  Rio-Grande-Valley-Lab-Ae-aegypti-49_S49_L001_R1_001 (paired).R2.fastq.gz | Lab | Exposed | RGV | Batch_1 |
| Rio-Grande-Valley-Lab-Ae-aegypti-50_S50 | Rio-Grande-Valley-Lab-Ae-aegypti-50_S50_L001_R1_001 (paired).R1.fastq.gz  Rio-Grande-Valley-Lab-Ae-aegypti-50_S50_L001_R1_001 (paired).R2.fastq.gz | Lab | Exposed | RGV | Batch_1 |
| Rio-Grande-Valley-Lab-Ae-aegypti-51_S51 | Rio-Grande-Valley-Lab-Ae-aegypti-51_S51_L001_R1_001 (paired).R1.fastq.gz  Rio-Grande-Valley-Lab-Ae-aegypti-51_S51_L001_R1_001 (paired).R2.fastq.gz | Lab | Exposed | RGV | Batch_1 |
| Rio-Grande-Valley-Lab-Ae-aegypti-52_S52 | Rio-Grande-Valley-Lab-Ae-aegypti-52_S52_L001_R1_001 (paired).R1.fastq.gz  Rio-Grande-Valley-Lab-Ae-aegypti-52_S52_L001_R1_001 (paired).R2.fastq.gz | Lab | Infected | RGV | Batch_1 |
| Rio-Grande-Valley-Lab-Ae-aegypti-53_S53 | Rio-Grande-Valley-Lab-Ae-aegypti-53_S53_L001_R1_001 (paired).R1.fastq.gz  Rio-Grande-Valley-Lab-Ae-aegypti-53_S53_L001_R1_001 (paired).R2.fastq.gz | Lab | Exposed | RGV | Batch_1 |
| Rio-Grande-Valley-Lab-Ae-aegypti-54_S54 | Rio-Grande-Valley-Lab-Ae-aegypti-54_S54_L001_R1_001 (paired).R1.fastq.gz  Rio-Grande-Valley-Lab-Ae-aegypti-54_S54_L001_R1_001 (paired).R2.fastq.gz | Lab | Infected | RGV | Batch_1 |
| Rio-Grande-Valley-Lab-Ae-aegypti-55_S55 | Rio-Grande-Valley-Lab-Ae-aegypti-55_S55_L001_R1_001 (paired).R1.fastq.gz  Rio-Grande-Valley-Lab-Ae-aegypti-55_S55_L001_R1_001 (paired).R2.fastq.gz | Lab | Exposed | RGV | Batch_1 |
| Rio-Grande-Valley-Lab-Ae-aegypti-56_S56 | Rio-Grande-Valley-Lab-Ae-aegypti-56_S56_L001_R1_001 (paired).R1.fastq.gz  Rio-Grande-Valley-Lab-Ae-aegypti-56_S56_L001_R1_001 (paired).R2.fastq.gz | Lab | Infected | RGV | Batch_1 |
| Rio-Grande-Valley-Lab-Ae-aegypti-57_S57 | Rio-Grande-Valley-Lab-Ae-aegypti-57_S57_L001_R1_001 (paired).R1.fastq.gz  Rio-Grande-Valley-Lab-Ae-aegypti-57_S57_L001_R1_001 (paired).R2.fastq.gz | Lab | Infected | RGV | Batch_1 |
| Rio-Grande-Valley-Lab-Ae-aegypti-58_S58 | Rio-Grande-Valley-Lab-Ae-aegypti-58_S58_L001_R1_001 (paired).R1.fastq.gz  Rio-Grande-Valley-Lab-Ae-aegypti-58_S58_L001_R1_001 (paired).R2.fastq.gz | Lab | Infected | RGV | Batch_1 |
| Rio-Grande-Valley-Lab-Ae-aegypti-59_S59 | Rio-Grande-Valley-Lab-Ae-aegypti-59_S59_L001_R1_001 (paired).R1.fastq.gz  Rio-Grande-Valley-Lab-Ae-aegypti-59_S59_L001_R1_001 (paired).R2.fastq.gz | Lab | Infected | RGV | Batch_1 |
| Rio-Grande-Valley-Lab-Ae-aegypti-60_S60 | Rio-Grande-Valley-Lab-Ae-aegypti-60_S60_L001_R1_001 (paired).R1.fastq.gz  Rio-Grande-Valley-Lab-Ae-aegypti-60_S60_L001_R1_001 (paired).R2.fastq.gz | Lab | Infected | RGV | Batch_1 |
| Rio-Grande-Valley-Lab-Ae-aegypti-61_S61 | Rio-Grande-Valley-Lab-Ae-aegypti-61_S61_L001_R1_001 (paired).R1.fastq.gz  Rio-Grande-Valley-Lab-Ae-aegypti-61_S61_L001_R1_001 (paired).R2.fastq.gz | Lab | Infected | RGV | Batch_1 |
| Rio-Grande-Valley-Lab-Ae-aegypti-62_S62 | Rio-Grande-Valley-Lab-Ae-aegypti-62_S62_L001_R1_001 (paired).R1.fastq.gz  Rio-Grande-Valley-Lab-Ae-aegypti-62_S62_L001_R1_001 (paired).R2.fastq.gz | Lab | Exposed | RGV | Batch_1 |
| Rio-Grande-Valley-Lab-Ae-aegypti-63_S63 | Rio-Grande-Valley-Lab-Ae-aegypti-63_S63_L001_R1_001 (paired).R1.fastq.gz  Rio-Grande-Valley-Lab-Ae-aegypti-63_S63_L001_R1_001 (paired).R2.fastq.gz | Lab | Infected | RGV | Batch_1 |
| Rio-Grande-Valley-Lab-Ae-aegypti-64_S64 | Rio-Grande-Valley-Lab-Ae-aegypti-64_S64_L001_R1_001 (paired).R1.fastq.gz  Rio-Grande-Valley-Lab-Ae-aegypti-64_S64_L001_R1_001 (paired).R2.fastq.gz | Lab | Exposed | RGV | Batch_1 |
| Rio-Grande-Valley-Lab-Ae-aegypti-65_S65 | Rio-Grande-Valley-Lab-Ae-aegypti-65_S65_L001_R1_001 (paired).R1.fastq.gz  Rio-Grande-Valley-Lab-Ae-aegypti-65_S65_L001_R1_001 (paired).R2.fastq.gz | Lab | Exposed | RGV | Batch_1 |
| Rio-Grande-Valley-Lab-Ae-aegypti-66_S66 | Rio-Grande-Valley-Lab-Ae-aegypti-66_S66_L001_R1_001 (paired).R1.fastq.gz  Rio-Grande-Valley-Lab-Ae-aegypti-66_S66_L001_R1_001 (paired).R2.fastq.gz | Lab | Exposed | RGV | Batch_1 |
| Rio-Grande-Valley-Lab-Ae-aegypti-67_S67 | Rio-Grande-Valley-Lab-Ae-aegypti-67_S67_L001_R1_001 (paired).R1.fastq.gz  Rio-Grande-Valley-Lab-Ae-aegypti-67_S67_L001_R1_001 (paired).R2.fastq.gz | Lab | Infected | RGV | Batch_1 |
| Rio-Grande-Valley-Lab-Ae-aegypti-68_S68 | Rio-Grande-Valley-Lab-Ae-aegypti-68_S68_L001_R1_001 (paired).R1.fastq.gz  Rio-Grande-Valley-Lab-Ae-aegypti-68_S68_L001_R1_001 (paired).R2.fastq.gz | Lab | Exposed | RGV | Batch_1 |
| Rio-Grande-Valley-Lab-Ae-aegypti-69_S69 | Rio-Grande-Valley-Lab-Ae-aegypti-69_S69_L001_R1_001 (paired).R1.fastq.gz  Rio-Grande-Valley-Lab-Ae-aegypti-69_S69_L001_R1_001 (paired).R2.fastq.gz | Lab | Exposed | RGV | Batch_1 |
| Rio-Grande-Valley-Lab-Ae-aegypti-70_S70 | Rio-Grande-Valley-Lab-Ae-aegypti-70_S70_L001_R1_001 (paired).R1.fastq.gz  Rio-Grande-Valley-Lab-Ae-aegypti-70_S70_L001_R1_001 (paired).R2.fastq.gz | Lab | Exposed | RGV | Batch_1 |
| Rio-Grande-Valley-Lab-Ae-aegypti-71_S71 | Rio-Grande-Valley-Lab-Ae-aegypti-71_S71_L001_R1_001 (paired).R1.fastq.gz  Rio-Grande-Valley-Lab-Ae-aegypti-71_S71_L001_R1_001 (paired).R2.fastq.gz | Lab | Exposed | RGV | Batch_1 |
| Rio-Grande-Valley-Lab-Ae-aegypti-72_S72 | Rio-Grande-Valley-Lab-Ae-aegypti-72_S72_L001_R1_001 (paired).R1.fastq.gz  Rio-Grande-Valley-Lab-Ae-aegypti-72_S72_L001_R1_001 (paired).R2.fastq.gz | Lab | Exposed | RGV | Batch_1 |
| Rio-Grande-Valley-Lab-Ae-aegypti-73_S73 | Rio-Grande-Valley-Lab-Ae-aegypti-73_S73_L001_R1_001 (paired).R1.fastq.gz  Rio-Grande-Valley-Lab-Ae-aegypti-73_S73_L001_R1_001 (paired).R2.fastq.gz | Lab | Infected | RGV | Batch_1 |
| Rio-Grande-Valley-Lab-Ae-aegypti-74_S74 | Rio-Grande-Valley-Lab-Ae-aegypti-74_S74_L001_R1_001 (paired).R1.fastq.gz  Rio-Grande-Valley-Lab-Ae-aegypti-74_S74_L001_R1_001 (paired).R2.fastq.gz | Lab | Infected | RGV | Batch_1 |
| Rio-Grande-Valley-Lab-Ae-aegypti-75_S75 | Rio-Grande-Valley-Lab-Ae-aegypti-75_S75_L001_R1_001 (paired).R1.fastq.gz  Rio-Grande-Valley-Lab-Ae-aegypti-75_S75_L001_R1_001 (paired).R2.fastq.gz | Lab | Infected | RGV | Batch_1 |
| Rio-Grande-Valley-Lab-Ae-aegypti-76_S76 | Rio-Grande-Valley-Lab-Ae-aegypti-76_S76_L001_R1_001 (paired).R1.fastq.gz  Rio-Grande-Valley-Lab-Ae-aegypti-76_S76_L001_R1_001 (paired).R2.fastq.gz | Lab | Infected | RGV | Batch_1 |
| Rio-Grande-Valley-Lab-Ae-aegypti-77_S77 | Rio-Grande-Valley-Lab-Ae-aegypti-77_S77_L001_R1_001 (paired).R1.fastq.gz  Rio-Grande-Valley-Lab-Ae-aegypti-77_S77_L001_R1_001 (paired).R2.fastq.gz | Lab | Infected | RGV | Batch_1 |
| Rio-Grande-Valley-Lab-Ae-aegypti-78_S78 | Rio-Grande-Valley-Lab-Ae-aegypti-78_S78_L001_R1_001 (paired).R1.fastq.gz  Rio-Grande-Valley-Lab-Ae-aegypti-78_S78_L001_R1_001 (paired).R2.fastq.gz | Lab | Infected | RGV | Batch_1 |
| Rio-Grande-Valley-Lab-Ae-aegypti-79_S79 | Rio-Grande-Valley-Lab-Ae-aegypti-79_S79_L001_R1_001 (paired).R1.fastq.gz  Rio-Grande-Valley-Lab-Ae-aegypti-79_S79_L001_R1_001 (paired).R2.fastq.gz | Lab | Exposed | RGV | Batch_1 |
| Rio-Grande-Valley-Lab-Ae-aegypti-80_S80 | Rio-Grande-Valley-Lab-Ae-aegypti-80_S80_L001_R1_001 (paired).R1.fastq.gz  Rio-Grande-Valley-Lab-Ae-aegypti-80_S80_L001_R1_001 (paired).R2.fastq.gz | Lab | Infected | RGV | Batch_1 |
| Rio-Grande-Valley-Lab-Ae-aegypti-81_S81 | Rio-Grande-Valley-Lab-Ae-aegypti-81_S81_L001_R1_001 (paired).R1.fastq.gz  Rio-Grande-Valley-Lab-Ae-aegypti-81_S81_L001_R1_001 (paired).R2.fastq.gz | Lab | Infected | RGV | Batch_1 |
| Rio-Grande-Valley-Lab-Ae-aegypti-82_S82 | Rio-Grande-Valley-Lab-Ae-aegypti-82_S82_L001_R1_001 (paired).R1.fastq.gz  Rio-Grande-Valley-Lab-Ae-aegypti-82_S82_L001_R1_001 (paired).R2.fastq.gz | Lab | Exposed | RGV | Batch_1 |
| Rio-Grande-Valley-Lab-Ae-aegypti-83_S83 | Rio-Grande-Valley-Lab-Ae-aegypti-83_S83_L001_R1_001 (paired).R1.fastq.gz  Rio-Grande-Valley-Lab-Ae-aegypti-83_S83_L001_R1_001 (paired).R2.fastq.gz | Lab | Infected | RGV | Batch_1 |
| Rio-Grande-Valley-Lab-Ae-aegypti-84_S84 | Rio-Grande-Valley-Lab-Ae-aegypti-84_S84_L001_R1_001 (paired).R1.fastq.gz  Rio-Grande-Valley-Lab-Ae-aegypti-84_S84_L001_R1_001 (paired).R2.fastq.gz | Lab | Infected | RGV | Batch_1 |
| Rio-Grande-Valley-Lab-Ae-aegypti-85_S85 | Rio-Grande-Valley-Lab-Ae-aegypti-85_S85_L001_R1_001 (paired).R1.fastq.gz  Rio-Grande-Valley-Lab-Ae-aegypti-85_S85_L001_R1_001 (paired).R2.fastq.gz | Lab | Exposed | RGV | Batch_1 |
| Rio-Grande-Valley-Lab-Ae-aegypti-Mock-1_S86 | Rio-Grande-Valley-Lab-Ae-aegypti-Mock-1_S86_L001_R1_001 (paired).R1.fastq.gz  Rio-Grande-Valley-Lab-Ae-aegypti-Mock-1_S86_L001_R1_001 (paired).R2.fastq.gz | Lab | Unexposed | RGV | Batch_1 |
| Rio-Grande-Valley-Lab-Ae-aegypti-Mock-2_S87 | Rio-Grande-Valley-Lab-Ae-aegypti-Mock-2_S87_L001_R1_001 (paired).R1.fastq.gz  Rio-Grande-Valley-Lab-Ae-aegypti-Mock-2_S87_L001_R1_001 (paired).R2.fastq.gz | Lab | Unexposed | RGV | Batch_1 |
| Rio-Grande-Valley-Lab-Ae-aegypti-Mock-3_S88 | Rio-Grande-Valley-Lab-Ae-aegypti-Mock-3_S88_L001_R1_001 (paired).R1.fastq.gz  Rio-Grande-Valley-Lab-Ae-aegypti-Mock-3_S88_L001_R1_001 (paired).R2.fastq.gz | Lab | Unexposed | RGV | Batch_1 |
| Rio-Grande-Valley-Lab-Ae-aegypti-Mock-4_S89 | Rio-Grande-Valley-Lab-Ae-aegypti-Mock-4_S89_L001_R1_001 (paired).R1.fastq.gz  Rio-Grande-Valley-Lab-Ae-aegypti-Mock-4_S89_L001_R1_001 (paired).R2.fastq.gz | Lab | Unexposed | RGV | Batch_1 |
| Rio-Grande-Valley-Lab-Ae-aegypti-Mock-5_S90 | Rio-Grande-Valley-Lab-Ae-aegypti-Mock-5_S90_L001_R1_001 (paired).R1.fastq.gz  Rio-Grande-Valley-Lab-Ae-aegypti-Mock-5_S90_L001_R1_001 (paired).R2.fastq.gz | Lab | Unexposed | RGV | Batch_1 |
| Rio-Grande-Valley-Lab-Ae-aegypti-Mock-6_S91 | Rio-Grande-Valley-Lab-Ae-aegypti-Mock-6_S91_L001_R1_001 (paired).R1.fastq.gz  Rio-Grande-Valley-Lab-Ae-aegypti-Mock-6_S91_L001_R1_001 (paired).R2.fastq.gz | Lab | Unexposed | RGV | Batch_1 |
| Rio-Grande-Valley-Lab-Ae-aegypti-Mock-7_S92 | Rio-Grande-Valley-Lab-Ae-aegypti-Mock-7_S92_L001_R1_001 (paired).R1.fastq.gz  Rio-Grande-Valley-Lab-Ae-aegypti-Mock-7_S92_L001_R1_001 (paired).R2.fastq.gz | Lab | Unexposed | RGV | Batch_1 |
| Rio-Grande-Valley-Lab-Ae-aegypti-Mock-8_S93 | Rio-Grande-Valley-Lab-Ae-aegypti-Mock-8_S93_L001_R1_001 (paired).R1.fastq.gz  Rio-Grande-Valley-Lab-Ae-aegypti-Mock-8_S93_L001_R1_001 (paired).R2.fastq.gz | Lab | Unexposed | RGV | Batch_1 |
| Rio-Grande-Valley-Lab-Ae-aegypti-Mock-9_S94 | Rio-Grande-Valley-Lab-Ae-aegypti-Mock-9_S94_L001_R1_001 (paired).R1.fastq.gz  Rio-Grande-Valley-Lab-Ae-aegypti-Mock-9_S94_L001_R1_001 (paired).R2.fastq.gz | Lab | Unexposed | RGV | Batch_1 |
| Rio-Grande-Valley-Lab-Ae-aegypti-Mock-10_S95 | Rio-Grande-Valley-Lab-Ae-aegypti-Mock-10_S95_L001_R1_001 (paired).R1.fastq.gz  Rio-Grande-Valley-Lab-Ae-aegypti-Mock-10_S95_L001_R1_001 (paired).R2.fastq.gz | Lab | Unexposed | RGV | Batch_1 |
| Rio-Grande-Valley-Lab-Ae-aegypti-Mock-11_S96 | Rio-Grande-Valley-Lab-Ae-aegypti-Mock-11_S96_L001_R1_001 (paired).R1.fastq.gz  Rio-Grande-Valley-Lab-Ae-aegypti-Mock-11_S96_L001_R1_001 (paired).R2.fastq.gz | Lab | Unexposed | RGV | Batch_1 |
| Rio-Grande-Valley-Lab-Ae-aegypti-Mock-13_S98 | Rio-Grande-Valley-Lab-Ae-aegypti-Mock-13_S98_L001_R1_001 (paired).R1.fastq.gz  Rio-Grande-Valley-Lab-Ae-aegypti-Mock-13_S98_L001_R1_001 (paired).R2.fastq.gz | Lab | Unexposed | RGV | Batch_1 |
| Rio-Grande-Valley-Lab-Ae-aegypti-Mock-14_S99 | Rio-Grande-Valley-Lab-Ae-aegypti-Mock-14_S99_L001_R1_001 (paired).R1.fastq.gz  Rio-Grande-Valley-Lab-Ae-aegypti-Mock-14_S99_L001_R1_001 (paired).R2.fastq.gz | Lab | Unexposed | RGV | Batch_1 |
| Rio-Grande-Valley-Lab-Ae-aegypti-Mock-15_S100 | Rio-Grande-Valley-Lab-Ae-aegypti-Mock-15_S100_L001_R1_001 (paired).R1.fastq.gz  Rio-Grande-Valley-Lab-Ae-aegypti-Mock-15_S100_L001_R1_001 (paired).R2.fastq.gz | Lab | Unexposed | RGV | Batch_1 |
| Rio-Grande-Valley-Lab-Ae-aegypti-Mock-16_S101 | Rio-Grande-Valley-Lab-Ae-aegypti-Mock-16_S101_L001_R1_001 (paired).R1.fastq.gz  Rio-Grande-Valley-Lab-Ae-aegypti-Mock-16_S101_L001_R1_001 (paired).R2.fastq.gz | Lab | Unexposed | RGV | Batch_1 |
| Rio-Grande-Valley-Lab-Ae-aegypti-Mock-17_S102 | Rio-Grande-Valley-Lab-Ae-aegypti-Mock-17_S102_L001_R1_001 (paired).R1.fastq.gz  Rio-Grande-Valley-Lab-Ae-aegypti-Mock-17_S102_L001_R1_001 (paired).R2.fastq.gz | Lab | Unexposed | RGV | Batch_1 |
| Rio-Grande-Valley-Lab-Ae-aegypti-Mock-18_S103 | Rio-Grande-Valley-Lab-Ae-aegypti-Mock-18_S103_L001_R1_001 (paired).R1.fastq.gz  Rio-Grande-Valley-Lab-Ae-aegypti-Mock-18_S103_L001_R1_001 (paired).R2.fastq.gz | Lab | Unexposed | RGV | Batch_1 |
| Rio-Grande-Valley-Lab-Ae-aegypti-Mock-19_S104 | Rio-Grande-Valley-Lab-Ae-aegypti-Mock-19_S104_L001_R1_001 (paired).R1.fastq.gz  Rio-Grande-Valley-Lab-Ae-aegypti-Mock-19_S104_L001_R1_001 (paired).R2.fastq.gz | Lab | Unexposed | RGV | Batch_1 |
| Rio-Grande-Valley-Lab-Ae-aegypti-Mock-20_S105 | Rio-Grande-Valley-Lab-Ae-aegypti-Mock-20_S105_L001_R1_001 (paired).R1.fastq.gz  Rio-Grande-Valley-Lab-Ae-aegypti-Mock-20_S105_L001_R1_001 (paired).R2.fastq.gz | Lab | Unexposed | RGV | Batch_1 |
| Rio-Grande-Valley-Lab-Ae-aegypti-Mock-21_S106 | Rio-Grande-Valley-Lab-Ae-aegypti-Mock-21_S106_L001_R1_001 (paired).R1.fastq.gz  Rio-Grande-Valley-Lab-Ae-aegypti-Mock-21_S106_L001_R1_001 (paired).R2.fastq.gz | Lab | Unexposed | RGV | Batch_1 |
| Rio-Grande-Valley-Lab-Ae-aegypti-Mock-22_S107 | Rio-Grande-Valley-Lab-Ae-aegypti-Mock-22_S107_L001_R1_001 (paired).R1.fastq.gz  Rio-Grande-Valley-Lab-Ae-aegypti-Mock-22_S107_L001_R1_001 (paired).R2.fastq.gz | Lab | Unexposed | RGV | Batch_1 |
| Rio-Grande-Valley-Lab-Ae-aegypti-Mock-23_S108 | Rio-Grande-Valley-Lab-Ae-aegypti-Mock-23_S108_L001_R1_001 (paired).R1.fastq.gz  Rio-Grande-Valley-Lab-Ae-aegypti-Mock-23_S108_L001_R1_001 (paired).R2.fastq.gz | Lab | Unexposed | RGV | Batch_1 |
| Rio-Grande-Valley-Lab-Ae-aegypti-Mock-24_S109 | Rio-Grande-Valley-Lab-Ae-aegypti-Mock-24_S109_L001_R1_001 (paired).R1.fastq.gz  Rio-Grande-Valley-Lab-Ae-aegypti-Mock-24_S109_L001_R1_001 (paired).R2.fastq.gz | Lab | Unexposed | RGV | Batch_1 |
| Rio-Grande-Valley-Lab-Ae-aegypti-Mock-25_S110 | Rio-Grande-Valley-Lab-Ae-aegypti-Mock-25_S110_L001_R1_001 (paired).R1.fastq.gz  Rio-Grande-Valley-Lab-Ae-aegypti-Mock-25_S110_L001_R1_001 (paired).R2.fastq.gz | Lab | Unexposed | RGV | Batch_1 |
| Rio-Grande-Valley-Lab-Ae-aegypti-Mock-26_S111 | Rio-Grande-Valley-Lab-Ae-aegypti-Mock-26_S111_L001_R1_001 (paired).R1.fastq.gz  Rio-Grande-Valley-Lab-Ae-aegypti-Mock-26_S111_L001_R1_001 (paired).R2.fastq.gz | Lab | Unexposed | RGV | Batch_1 |
| Rio-Grande-Valley-Lab-Ae-aegypti-Mock-27_S112 | Rio-Grande-Valley-Lab-Ae-aegypti-Mock-27_S112_L001_R1_001 (paired).R1.fastq.gz  Rio-Grande-Valley-Lab-Ae-aegypti-Mock-27_S112_L001_R1_001 (paired).R2.fastq.gz | Lab | Unexposed | RGV | Batch_1 |
| Rio-Grande-Valley-Lab-Ae-aegypti-Mock-28_S113 | Rio-Grande-Valley-Lab-Ae-aegypti-Mock-28_S113_L001_R1_001 (paired).R1.fastq.gz  Rio-Grande-Valley-Lab-Ae-aegypti-Mock-28_S113_L001_R1_001 (paired).R2.fastq.gz | Lab | Unexposed | RGV | Batch_1 |
| Rio-Grande-Valley-Lab-Ae-aegypti-Mock-29_S114 | Rio-Grande-Valley-Lab-Ae-aegypti-Mock-29_S114_L001_R1_001 (paired).R1.fastq.gz  Rio-Grande-Valley-Lab-Ae-aegypti-Mock-29_S114_L001_R1_001 (paired).R2.fastq.gz | Lab | Unexposed | RGV | Batch_1 |
| Rio-Grande-Valley-Lab-Ae-aegypti-Mock-30_S115 | Rio-Grande-Valley-Lab-Ae-aegypti-Mock-30_S115_L001_R1_001 (paired).R1.fastq.gz  Rio-Grande-Valley-Lab-Ae-aegypti-Mock-30_S115_L001_R1_001 (paired).R2.fastq.gz | Lab | Unexposed | RGV | Batch_1 |
| Rio-Grande-Valley-Lab-Ae-aegypti-Mock-31_S116 | Rio-Grande-Valley-Lab-Ae-aegypti-Mock-31_S116_L001_R1_001 (paired).R1.fastq.gz  Rio-Grande-Valley-Lab-Ae-aegypti-Mock-31_S116_L001_R1_001 (paired).R2.fastq.gz | Lab | Unexposed | RGV | Batch_1 |
| Rio-Grande-Valley-Lab-Ae-aegypti-Mock-32_S117 | Rio-Grande-Valley-Lab-Ae-aegypti-Mock-32_S117_L001_R1_001 (paired).R1.fastq.gz  Rio-Grande-Valley-Lab-Ae-aegypti-Mock-32_S117_L001_R1_001 (paired).R2.fastq.gz | Lab | Unexposed | RGV | Batch_1 |
| Rio-Grande-Valley-Lab-Ae-aegypti-Mock-33_S118 | Rio-Grande-Valley-Lab-Ae-aegypti-Mock-33_S118_L001_R1_001 (paired).R1.fastq.gz  Rio-Grande-Valley-Lab-Ae-aegypti-Mock-33_S118_L001_R1_001 (paired).R2.fastq.gz | Lab | Unexposed | RGV | Batch_1 |
| Rio-Grande-Valley-Lab-Ae-aegypti-Mock-34_S119 | Rio-Grande-Valley-Lab-Ae-aegypti-Mock-34_S119_L001_R1_001 (paired).R1.fastq.gz  Rio-Grande-Valley-Lab-Ae-aegypti-Mock-34_S119_L001_R1_001 (paired).R2.fastq.gz | Lab | Unexposed | RGV | Batch_1 |
| Rio-Grande-Valley-Lab-Ae-aegypti-Mock-36_S121 | Rio-Grande-Valley-Lab-Ae-aegypti-Mock-36_S121_L001_R1_001 (paired).R1.fastq.gz  Rio-Grande-Valley-Lab-Ae-aegypti-Mock-36_S121_L001_R1_001 (paired).R2.fastq.gz | Lab | Unexposed | RGV | Batch_1 |
| Rio-Grande-Valley-Lab-Ae-aegypti-Mock-37_S122 | Rio-Grande-Valley-Lab-Ae-aegypti-Mock-37_S122_L001_R1_001 (paired).R1.fastq.gz  Rio-Grande-Valley-Lab-Ae-aegypti-Mock-37_S122_L001_R1_001 (paired).R2.fastq.gz | Lab | Unexposed | RGV | Batch_1 |
| Rio-Grande-Valley-Lab-Ae-aegypti-Mock-38_S123 | Rio-Grande-Valley-Lab-Ae-aegypti-Mock-38_S123_L001_R1_001 (paired).R1.fastq.gz  Rio-Grande-Valley-Lab-Ae-aegypti-Mock-38_S123_L001_R1_001 (paired).R2.fastq.gz | Lab | Unexposed | RGV | Batch_1 |
| Rio-Grande-Valley-Lab-Ae-aegypti-Mock-39_S124 | Rio-Grande-Valley-Lab-Ae-aegypti-Mock-39_S124_L001_R1_001 (paired).R1.fastq.gz  Rio-Grande-Valley-Lab-Ae-aegypti-Mock-39_S124_L001_R1_001 (paired).R2.fastq.gz | Lab | Unexposed | RGV | Batch_1 |
| Rio-Grande-Valley-Lab-Ae-aegypti-Mock-40_S125 | Rio-Grande-Valley-Lab-Ae-aegypti-Mock-40_S125_L001_R1_001 (paired).R1.fastq.gz  Rio-Grande-Valley-Lab-Ae-aegypti-Mock-40_S125_L001_R1_001 (paired).R2.fastq.gz | Lab | Unexposed | RGV | Batch_1 |
| Galveston-Ae-aegypti-Lab-1_S126 | Galveston-Ae-aegypti-Lab-1_S126_L001_R1_001 (paired).R1.fastq.gz  Galveston-Ae-aegypti-Lab-1_S126_L001_R1_001 (paired).R2.fastq.gz | Lab | Infected | Galveston | Batch_1 |
| Galveston-Ae-aegypti-Lab-2_S127 | Galveston-Ae-aegypti-Lab-2_S127_L001_R1_001 (paired).R1.fastq.gz  Galveston-Ae-aegypti-Lab-2_S127_L001_R1_001 (paired).R2.fastq.gz | Lab | Exposed | Galveston | Batch_1 |
| Galveston-Ae-aegypti-Lab-3_S128 | Galveston-Ae-aegypti-Lab-3_S128_L001_R1_001 (paired).R1.fastq.gz  Galveston-Ae-aegypti-Lab-3_S128_L001_R1_001 (paired).R2.fastq.gz | Lab | Infected | Galveston | Batch_1 |
| Galveston-Ae-aegypti-Lab-4_S129 | Galveston-Ae-aegypti-Lab-4_S129_L001_R1_001 (paired).R1.fastq.gz  Galveston-Ae-aegypti-Lab-4_S129_L001_R1_001 (paired).R2.fastq.gz | Lab | Infected | Galveston | Batch_1 |
| Galveston-Ae-aegypti-Lab-5_S130 | Galveston-Ae-aegypti-Lab-5_S130_L001_R1_001 (paired).R1.fastq.gz  Galveston-Ae-aegypti-Lab-5_S130_L001_R1_001 (paired).R2.fastq.gz | Lab | Infected | Galveston | Batch_1 |
| Galveston-Ae-aegypti-Lab-6_S131 | Galveston-Ae-aegypti-Lab-6_S131_L001_R1_001 (paired).R1.fastq.gz  Galveston-Ae-aegypti-Lab-6_S131_L001_R1_001 (paired).R2.fastq.gz | Lab | Exposed | Galveston | Batch_1 |
| Galveston-Ae-aegypti-Lab-7_S132 | Galveston-Ae-aegypti-Lab-7_S132_L001_R1_001 (paired).R1.fastq.gz  Galveston-Ae-aegypti-Lab-7_S132_L001_R1_001 (paired).R2.fastq.gz | Lab | Infected | Galveston | Batch_1 |
| Galveston-Ae-aegypti-Lab-8_S133 | Galveston-Ae-aegypti-Lab-8_S133_L001_R1_001 (paired).R1.fastq.gz  Galveston-Ae-aegypti-Lab-8_S133_L001_R1_001 (paired).R2.fastq.gz | Lab | Infected | Galveston | Batch_1 |
| Galveston-Ae-aegypti-Lab-9_S134 | Galveston-Ae-aegypti-Lab-9_S134_L001_R1_001 (paired).R1.fastq.gz  Galveston-Ae-aegypti-Lab-9_S134_L001_R1_001 (paired).R2.fastq.gz | Lab | Infected | Galveston | Batch_1 |
| Galveston-Ae-aegypti-Lab-10_S135 | Galveston-Ae-aegypti-Lab-10_S135_L001_R1_001 (paired).R1.fastq.gz  Galveston-Ae-aegypti-Lab-10_S135_L001_R1_001 (paired).R2.fastq.gz | Lab | Infected | Galveston | Batch_1 |
| Galveston-Ae-aegypti-Lab-11_S136 | Galveston-Ae-aegypti-Lab-11_S136_L001_R1_001 (paired).R1.fastq.gz  Galveston-Ae-aegypti-Lab-11_S136_L001_R1_001 (paired).R2.fastq.gz | Lab | Infected | Galveston | Batch_1 |
| Galveston-Ae-aegypti-Lab-12_S137 | Galveston-Ae-aegypti-Lab-12_S137_L001_R1_001 (paired).R1.fastq.gz  Galveston-Ae-aegypti-Lab-12_S137_L001_R1_001 (paired).R2.fastq.gz | Lab | Infected | Galveston | Batch_1 |
| Galveston-Ae-aegypti-Lab-13_S138 | Galveston-Ae-aegypti-Lab-13_S138_L001_R1_001 (paired).R1.fastq.gz  Galveston-Ae-aegypti-Lab-13_S138_L001_R1_001 (paired).R2.fastq.gz | Lab | Infected | Galveston | Batch_1 |
| Galveston-Ae-aegypti-Lab-14_S139 | Galveston-Ae-aegypti-Lab-14_S139_L001_R1_001 (paired).R1.fastq.gz  Galveston-Ae-aegypti-Lab-14_S139_L001_R1_001 (paired).R2.fastq.gz | Lab | Exposed | Galveston | Batch_1 |
| Galveston-Ae-aegypti-Lab-15_S140 | Galveston-Ae-aegypti-Lab-15_S140_L001_R1_001 (paired).R1.fastq.gz  Galveston-Ae-aegypti-Lab-15_S140_L001_R1_001 (paired).R2.fastq.gz | Lab | Infected | Galveston | Batch_1 |
| Galveston-Ae-aegypti-Lab-16_S141 | Galveston-Ae-aegypti-Lab-16_S141_L001_R1_001 (paired).R1.fastq.gz  Galveston-Ae-aegypti-Lab-16_S141_L001_R1_001 (paired).R2.fastq.gz | Lab | Exposed | Galveston | Batch_1 |
| Galveston-Ae-aegypti-Lab-17_S142 | Galveston-Ae-aegypti-Lab-17_S142_L001_R1_001 (paired).R1.fastq.gz  Galveston-Ae-aegypti-Lab-17_S142_L001_R1_001 (paired).R2.fastq.gz | Lab | Infected | Galveston | Batch_1 |
| Galveston-Ae-aegypti-Lab-18_S143 | Galveston-Ae-aegypti-Lab-18_S143_L001_R1_001 (paired).R1.fastq.gz  Galveston-Ae-aegypti-Lab-18_S143_L001_R1_001 (paired).R2.fastq.gz | Lab | Exposed | Galveston | Batch_1 |
| Galveston-Ae-aegypti-Lab-19_S144 | Galveston-Ae-aegypti-Lab-19_S144_L001_R1_001 (paired).R1.fastq.gz  Galveston-Ae-aegypti-Lab-19_S144_L001_R1_001 (paired).R2.fastq.gz | Lab | Exposed | Galveston | Batch_1 |
| Galveston-Ae-aegypti-Lab-20_S145 | Galveston-Ae-aegypti-Lab-20_S145_L001_R1_001 (paired).R1.fastq.gz  Galveston-Ae-aegypti-Lab-20_S145_L001_R1_001 (paired).R2.fastq.gz | Lab | Exposed | Galveston | Batch_1 |
| Galveston-Ae-aegypti-Lab-21_S146 | Galveston-Ae-aegypti-Lab-21_S146_L001_R1_001 (paired).R1.fastq.gz  Galveston-Ae-aegypti-Lab-21_S146_L001_R1_001 (paired).R2.fastq.gz | Lab | Infected | Galveston | Batch_1 |
| Galveston-Ae-aegypti-Lab-22_S147 | Galveston-Ae-aegypti-Lab-22_S147_L001_R1_001 (paired).R1.fastq.gz  Galveston-Ae-aegypti-Lab-22_S147_L001_R1_001 (paired).R2.fastq.gz | Lab | Exposed | Galveston | Batch_1 |
| Galveston-Ae-aegypti-Lab-23_S148 | Galveston-Ae-aegypti-Lab-23_S148_L001_R1_001 (paired).R1.fastq.gz  Galveston-Ae-aegypti-Lab-23_S148_L001_R1_001 (paired).R2.fastq.gz | Lab | Exposed | Galveston | Batch_1 |
| Galveston-Ae-aegypti-Lab-24_S149 | Galveston-Ae-aegypti-Lab-24_S149_L001_R1_001 (paired).R1.fastq.gz  Galveston-Ae-aegypti-Lab-24_S149_L001_R1_001 (paired).R2.fastq.gz | Lab | Infected | Galveston | Batch_1 |
| Galveston-Ae-aegypti-Lab-25_S150 | Galveston-Ae-aegypti-Lab-25_S150_L001_R1_001 (paired).R1.fastq.gz  Galveston-Ae-aegypti-Lab-25_S150_L001_R1_001 (paired).R2.fastq.gz | Lab | Infected | Galveston | Batch_1 |
| Galveston-Ae-aegypti-Lab-26_S151 | Galveston-Ae-aegypti-Lab-26_S151_L001_R1_001 (paired).R1.fastq.gz  Galveston-Ae-aegypti-Lab-26_S151_L001_R1_001 (paired).R2.fastq.gz | Lab | Exposed | Galveston | Batch_1 |
| Galveston-Ae-aegypti-Lab-27_S152 | Galveston-Ae-aegypti-Lab-27_S152_L001_R1_001 (paired).R1.fastq.gz  Galveston-Ae-aegypti-Lab-27_S152_L001_R1_001 (paired).R2.fastq.gz | Lab | Infected | Galveston | Batch_1 |
| Galveston-Ae-aegypti-Lab-28_S153 | Galveston-Ae-aegypti-Lab-28_S153_L001_R1_001 (paired).R1.fastq.gz  Galveston-Ae-aegypti-Lab-28_S153_L001_R1_001 (paired).R2.fastq.gz | Lab | Infected | Galveston | Batch_1 |
| Galveston-Ae-aegypti-Lab-29_S154 | Galveston-Ae-aegypti-Lab-29_S154_L001_R1_001 (paired).R1.fastq.gz  Galveston-Ae-aegypti-Lab-29_S154_L001_R1_001 (paired).R2.fastq.gz | Lab | Infected | Galveston | Batch_1 |
| Galveston-Ae-aegypti-Lab-30_S155 | Galveston-Ae-aegypti-Lab-30_S155_L001_R1_001 (paired).R1.fastq.gz  Galveston-Ae-aegypti-Lab-30_S155_L001_R1_001 (paired).R2.fastq.gz | Lab | Exposed | Galveston | Batch_1 |
| Galveston-Ae-aegypti-Lab-31_S156 | Galveston-Ae-aegypti-Lab-31_S156_L001_R1_001 (paired).R1.fastq.gz  Galveston-Ae-aegypti-Lab-31_S156_L001_R1_001 (paired).R2.fastq.gz | Lab | Exposed | Galveston | Batch_1 |
| Galveston-Ae-aegypti-Lab-32_S157 | Galveston-Ae-aegypti-Lab-32_S157_L001_R1_001 (paired).R1.fastq.gz  Galveston-Ae-aegypti-Lab-32_S157_L001_R1_001 (paired).R2.fastq.gz | Lab | Infected | Galveston | Batch_1 |
| Galveston-Ae-aegypti-Lab-33_S158 | Galveston-Ae-aegypti-Lab-33_S158_L001_R1_001 (paired).R1.fastq.gz  Galveston-Ae-aegypti-Lab-33_S158_L001_R1_001 (paired).R2.fastq.gz | Lab | Infected | Galveston | Batch_1 |
| Galveston-Ae-aegypti-Lab-34_S159 | Galveston-Ae-aegypti-Lab-34_S159_L001_R1_001 (paired).R1.fastq.gz  Galveston-Ae-aegypti-Lab-34_S159_L001_R1_001 (paired).R2.fastq.gz | Lab | Infected | Galveston | Batch_1 |
| Galveston-Ae-aegypti-Lab-35_S160 | Galveston-Ae-aegypti-Lab-35_S160_L001_R1_001 (paired).R1.fastq.gz  Galveston-Ae-aegypti-Lab-35_S160_L001_R1_001 (paired).R2.fastq.gz | Lab | Infected | Galveston | Batch_1 |
| Galveston-Ae-aegypti-Lab-36_S161 | Galveston-Ae-aegypti-Lab-36_S161_L001_R1_001 (paired).R1.fastq.gz  Galveston-Ae-aegypti-Lab-36_S161_L001_R1_001 (paired).R2.fastq.gz | Lab | Infected | Galveston | Batch_1 |
| Galveston-Ae-aegypti-Lab-37_S162 | Galveston-Ae-aegypti-Lab-37_S162_L001_R1_001 (paired).R1.fastq.gz  Galveston-Ae-aegypti-Lab-37_S162_L001_R1_001 (paired).R2.fastq.gz | Lab | Infected | Galveston | Batch_1 |
| Galveston-Ae-aegypti-Lab-38_S163 | Galveston-Ae-aegypti-Lab-38_S163_L001_R1_001 (paired).R1.fastq.gz  Galveston-Ae-aegypti-Lab-38_S163_L001_R1_001 (paired).R2.fastq.gz | Lab | Infected | Galveston | Batch_1 |
| Galveston-Ae-aegypti-Lab-39_S164 | Galveston-Ae-aegypti-Lab-39_S164_L001_R1_001 (paired).R1.fastq.gz  Galveston-Ae-aegypti-Lab-39_S164_L001_R1_001 (paired).R2.fastq.gz | Lab | Infected | Galveston | Batch_1 |
| Galveston-Ae-aegypti-Lab-40_S165 | Galveston-Ae-aegypti-Lab-40_S165_L001_R1_001 (paired).R1.fastq.gz  Galveston-Ae-aegypti-Lab-40_S165_L001_R1_001 (paired).R2.fastq.gz | Lab | Infected | Galveston | Batch_1 |
| Galveston-Ae-aegypti-Lab-41_S166 | Galveston-Ae-aegypti-Lab-41_S166_L001_R1_001 (paired).R1.fastq.gz  Galveston-Ae-aegypti-Lab-41_S166_L001_R1_001 (paired).R2.fastq.gz | Lab | Exposed | Galveston | Batch_1 |
| Galveston-Ae-aegypti-Lab-42_S167 | Galveston-Ae-aegypti-Lab-42_S167_L001_R1_001 (paired).R1.fastq.gz  Galveston-Ae-aegypti-Lab-42_S167_L001_R1_001 (paired).R2.fastq.gz | Lab | Infected | Galveston | Batch_1 |
| Galveston-Ae-aegypti-Lab-43_S168 | Galveston-Ae-aegypti-Lab-43_S168_L001_R1_001 (paired).R1.fastq.gz  Galveston-Ae-aegypti-Lab-43_S168_L001_R1_001 (paired).R2.fastq.gz | Lab | Infected | Galveston | Batch_1 |
| Galveston-Ae-aegypti-Lab-44_S169 | Galveston-Ae-aegypti-Lab-44_S169_L001_R1_001 (paired).R1.fastq.gz  Galveston-Ae-aegypti-Lab-44_S169_L001_R1_001 (paired).R2.fastq.gz | Lab | Infected | Galveston | Batch_1 |
| Galveston-Ae-aegypti-Lab-45_S170 | Galveston-Ae-aegypti-Lab-45_S170_L001_R1_001 (paired).R1.fastq.gz  Galveston-Ae-aegypti-Lab-45_S170_L001_R1_001 (paired).R2.fastq.gz | Lab | Infected | Galveston | Batch_1 |
| Galveston-Ae-aegypti-Lab-46_S171 | Galveston-Ae-aegypti-Lab-46_S171_L001_R1_001 (paired).R1.fastq.gz  Galveston-Ae-aegypti-Lab-46_S171_L001_R1_001 (paired).R2.fastq.gz | Lab | Infected | Galveston | Batch_1 |
| Galveston-Ae-aegypti-Lab-47_S172 | Galveston-Ae-aegypti-Lab-47_S172_L001_R1_001 (paired).R1.fastq.gz  Galveston-Ae-aegypti-Lab-47_S172_L001_R1_001 (paired).R2.fastq.gz | Lab | Infected | Galveston | Batch_1 |
| Galveston-Ae-aegypti-Lab-48_S173 | Galveston-Ae-aegypti-Lab-48_S173_L001_R1_001 (paired).R1.fastq.gz  Galveston-Ae-aegypti-Lab-48_S173_L001_R1_001 (paired).R2.fastq.gz | Lab | Exposed | Galveston | Batch_1 |
| Galveston-Ae-aegypti-Lab-49_S174 | Galveston-Ae-aegypti-Lab-49_S174_L001_R1_001 (paired).R1.fastq.gz  Galveston-Ae-aegypti-Lab-49_S174_L001_R1_001 (paired).R2.fastq.gz | Lab | Infected | Galveston | Batch_1 |
| Galveston-Ae-aegypti-Lab-50_S175 | Galveston-Ae-aegypti-Lab-50_S175_L001_R1_001 (paired).R1.fastq.gz  Galveston-Ae-aegypti-Lab-50_S175_L001_R1_001 (paired).R2.fastq.gz | Lab | Infected | Galveston | Batch_1 |
| Galveston-Ae-aegypti-Lab-51_S176 | Galveston-Ae-aegypti-Lab-51_S176_L001_R1_001 (paired).R1.fastq.gz  Galveston-Ae-aegypti-Lab-51_S176_L001_R1_001 (paired).R2.fastq.gz | Lab | Infected | Galveston | Batch_1 |
| Galveston-Ae-aegypti-Lab-52_S177 | Galveston-Ae-aegypti-Lab-52_S177_L001_R1_001 (paired).R1.fastq.gz  Galveston-Ae-aegypti-Lab-52_S177_L001_R1_001 (paired).R2.fastq.gz | Lab | Infected | Galveston | Batch_1 |
| Galveston-Ae-aegypti-Lab-53_S178 | Galveston-Ae-aegypti-Lab-53_S178_L001_R1_001 (paired).R1.fastq.gz  Galveston-Ae-aegypti-Lab-53_S178_L001_R1_001 (paired).R2.fastq.gz | Lab | Exposed | Galveston | Batch_1 |
| Galveston-Ae-aegypti-Lab-54_S179 | Galveston-Ae-aegypti-Lab-54_S179_L001_R1_001 (paired).R1.fastq.gz  Galveston-Ae-aegypti-Lab-54_S179_L001_R1_001 (paired).R2.fastq.gz | Lab | Infected | Galveston | Batch_1 |
| Galveston-Ae-aegypti-Lab-55_S180 | Galveston-Ae-aegypti-Lab-55_S180_L001_R1_001 (paired).R1.fastq.gz  Galveston-Ae-aegypti-Lab-55_S180_L001_R1_001 (paired).R2.fastq.gz | Lab | Infected | Galveston | Batch_1 |
| Galveston-Ae-aegypti-Lab-56_S181 | Galveston-Ae-aegypti-Lab-56_S181_L001_R1_001 (paired).R1.fastq.gz  Galveston-Ae-aegypti-Lab-56_S181_L001_R1_001 (paired).R2.fastq.gz | Lab | Infected | Galveston | Batch_1 |
| Galveston-Ae-aegypti-Lab-57_S182 | Galveston-Ae-aegypti-Lab-57_S182_L001_R1_001 (paired).R1.fastq.gz  Galveston-Ae-aegypti-Lab-57_S182_L001_R1_001 (paired).R2.fastq.gz | Lab | Infected | Galveston | Batch_1 |
| Galveston-Ae-aegypti-Lab-Mock-1_S183 | Galveston-Ae-aegypti-Lab-Mock-1_S183_L001_R1_001 (paired).R1.fastq.gz  Galveston-Ae-aegypti-Lab-Mock-1_S183_L001_R1_001 (paired).R2.fastq.gz | Lab | Unexposed | Galveston | Batch_1 |
| Galveston-Ae-aegypti-Lab-Mock-2_S184 | Galveston-Ae-aegypti-Lab-Mock-2_S184_L001_R1_001 (paired).R1.fastq.gz  Galveston-Ae-aegypti-Lab-Mock-2_S184_L001_R1_001 (paired).R2.fastq.gz | Lab | Unexposed | Galveston | Batch_1 |
| Galveston-Ae-aegypti-Lab-Mock-3_S185 | Galveston-Ae-aegypti-Lab-Mock-3_S185_L001_R1_001 (paired).R1.fastq.gz  Galveston-Ae-aegypti-Lab-Mock-3_S185_L001_R1_001 (paired).R2.fastq.gz | Lab | Unexposed | Galveston | Batch_1 |
| Galveston-Ae-aegypti-Lab-Mock-4_S186 | Galveston-Ae-aegypti-Lab-Mock-4_S186_L001_R1_001 (paired).R1.fastq.gz  Galveston-Ae-aegypti-Lab-Mock-4_S186_L001_R1_001 (paired).R2.fastq.gz | Lab | Unexposed | Galveston | Batch_1 |
| Galveston-Ae-aegypti-Lab-Mock-5_S187 | Galveston-Ae-aegypti-Lab-Mock-5_S187_L001_R1_001 (paired).R1.fastq.gz  Galveston-Ae-aegypti-Lab-Mock-5_S187_L001_R1_001 (paired).R2.fastq.gz | Lab | Unexposed | Galveston | Batch_1 |
| Galveston-Ae-aegypti-Lab-Mock-6_S188 | Galveston-Ae-aegypti-Lab-Mock-6_S188_L001_R1_001 (paired).R1.fastq.gz  Galveston-Ae-aegypti-Lab-Mock-6_S188_L001_R1_001 (paired).R2.fastq.gz | Lab | Unexposed | Galveston | Batch_1 |
| Galveston-Ae-aegypti-Lab-Mock-7_S189 | Galveston-Ae-aegypti-Lab-Mock-7_S189_L001_R1_001 (paired).R1.fastq.gz  Galveston-Ae-aegypti-Lab-Mock-7_S189_L001_R1_001 (paired).R2.fastq.gz | Lab | Unexposed | Galveston | Batch_1 |
| Galveston-Ae-aegypti-Lab-Mock-8_S190 | Galveston-Ae-aegypti-Lab-Mock-8_S190_L001_R1_001 (paired).R1.fastq.gz  Galveston-Ae-aegypti-Lab-Mock-8_S190_L001_R1_001 (paired).R2.fastq.gz | Lab | Unexposed | Galveston | Batch_1 |
| Galveston-Ae-aegypti-Lab-Mock-9_S191 | Galveston-Ae-aegypti-Lab-Mock-9_S191_L001_R1_001 (paired).R1.fastq.gz  Galveston-Ae-aegypti-Lab-Mock-9_S191_L001_R1_001 (paired).R2.fastq.gz | Lab | Unexposed | Galveston | Batch_1 |
| Galveston-Ae-aegypti-Lab-Mock-10_S192 | Galveston-Ae-aegypti-Lab-Mock-10_S192_L001_R1_001 (paired).R1.fastq.gz  Galveston-Ae-aegypti-Lab-Mock-10_S192_L001_R1_001 (paired).R2.fastq.gz | Lab | Unexposed | Galveston | Batch_1 |
| Galveston-Ae-aegypti-Lab-Mock-11_S193 | Galveston-Ae-aegypti-Lab-Mock-11_S193_L001_R1_001 (paired).R1.fastq.gz  Galveston-Ae-aegypti-Lab-Mock-11_S193_L001_R1_001 (paired).R2.fastq.gz | Lab | Unexposed | Galveston | Batch_1 |
| Galveston-Ae-aegypti-Lab-Mock-12_S194 | Galveston-Ae-aegypti-Lab-Mock-12_S194_L001_R1_001 (paired).R1.fastq.gz  Galveston-Ae-aegypti-Lab-Mock-12_S194_L001_R1_001 (paired).R2.fastq.gz | Lab | Unexposed | Galveston | Batch_1 |
| Galveston-Ae-aegypti-Lab-Mock-13_S195 | Galveston-Ae-aegypti-Lab-Mock-13_S195_L001_R1_001 (paired).R1.fastq.gz  Galveston-Ae-aegypti-Lab-Mock-13_S195_L001_R1_001 (paired).R2.fastq.gz | Lab | Unexposed | Galveston | Batch_1 |
| Galveston-Ae-aegypti-Lab-Mock-14_S196 | Galveston-Ae-aegypti-Lab-Mock-14_S196_L001_R1_001 (paired).R1.fastq.gz  Galveston-Ae-aegypti-Lab-Mock-14_S196_L001_R1_001 (paired).R2.fastq.gz | Lab | Unexposed | Galveston | Batch_1 |
| Galveston-Ae-aegypti-Lab-Mock-15_S197 | Galveston-Ae-aegypti-Lab-Mock-15_S197_L001_R1_001 (paired).R1.fastq.gz  Galveston-Ae-aegypti-Lab-Mock-15_S197_L001_R1_001 (paired).R2.fastq.gz | Lab | Unexposed | Galveston | Batch_1 |
| Galveston-Ae-aegypti-Lab-Mock-16_S198 | Galveston-Ae-aegypti-Lab-Mock-16_S198_L001_R1_001 (paired).R1.fastq.gz  Galveston-Ae-aegypti-Lab-Mock-16_S198_L001_R1_001 (paired).R2.fastq.gz | Lab | Unexposed | Galveston | Batch_1 |
| Galveston-Ae-aegypti-Lab-Mock-17_S199 | Galveston-Ae-aegypti-Lab-Mock-17_S199_L001_R1_001 (paired).R1.fastq.gz  Galveston-Ae-aegypti-Lab-Mock-17_S199_L001_R1_001 (paired).R2.fastq.gz | Lab | Unexposed | Galveston | Batch_1 |
| Galveston-Ae-aegypti-Lab-Mock-18_S200 | Galveston-Ae-aegypti-Lab-Mock-18_S200_L001_R1_001 (paired).R1.fastq.gz  Galveston-Ae-aegypti-Lab-Mock-18_S200_L001_R1_001 (paired).R2.fastq.gz | Lab | Unexposed | Galveston | Batch_1 |
| Galveston-Ae-aegypti-Lab-Mock-19_S201 | Galveston-Ae-aegypti-Lab-Mock-19_S201_L001_R1_001 (paired).R1.fastq.gz  Galveston-Ae-aegypti-Lab-Mock-19_S201_L001_R1_001 (paired).R2.fastq.gz | Lab | Unexposed | Galveston | Batch_1 |
| Galveston-Ae-aegypti-Lab-Mock-20_S202 | Galveston-Ae-aegypti-Lab-Mock-20_S202_L001_R1_001 (paired).R1.fastq.gz  Galveston-Ae-aegypti-Lab-Mock-20_S202_L001_R1_001 (paired).R2.fastq.gz | Lab | Unexposed | Galveston | Batch_1 |
| Galveston-Ae-aegypti-Lab-Mock-21_S203 | Galveston-Ae-aegypti-Lab-Mock-21_S203_L001_R1_001 (paired).R1.fastq.gz  Galveston-Ae-aegypti-Lab-Mock-21_S203_L001_R1_001 (paired).R2.fastq.gz | Lab | Unexposed | Galveston | Batch_1 |
| Galveston-Ae-aegypti-Lab-Mock-22_S204 | Galveston-Ae-aegypti-Lab-Mock-22_S204_L001_R1_001 (paired).R1.fastq.gz  Galveston-Ae-aegypti-Lab-Mock-22_S204_L001_R1_001 (paired).R2.fastq.gz | Lab | Unexposed | Galveston | Batch_1 |
| Galveston-Ae-aegypti-Lab-Mock-23_S205 | Galveston-Ae-aegypti-Lab-Mock-23_S205_L001_R1_001 (paired).R1.fastq.gz  Galveston-Ae-aegypti-Lab-Mock-23_S205_L001_R1_001 (paired).R2.fastq.gz | Lab | Unexposed | Galveston | Batch_1 |
| Galveston-Ae-aegypti-Lab-Mock-24_S206 | Galveston-Ae-aegypti-Lab-Mock-24_S206_L001_R1_001 (paired).R1.fastq.gz  Galveston-Ae-aegypti-Lab-Mock-24_S206_L001_R1_001 (paired).R2.fastq.gz | Lab | Unexposed | Galveston | Batch_1 |
| Galveston-Ae-aegypti-Lab-Mock-25_S207 | Galveston-Ae-aegypti-Lab-Mock-25_S207_L001_R1_001 (paired).R1.fastq.gz  Galveston-Ae-aegypti-Lab-Mock-25_S207_L001_R1_001 (paired).R2.fastq.gz | Lab | Unexposed | Galveston | Batch_1 |
| Galveston-Ae-aegypti-Lab-Mock-26_S208 | Galveston-Ae-aegypti-Lab-Mock-26_S208_L001_R1_001 (paired).R1.fastq.gz  Galveston-Ae-aegypti-Lab-Mock-26_S208_L001_R1_001 (paired).R2.fastq.gz | Lab | Unexposed | Galveston | Batch_1 |
| Galveston-Ae-aegypti-Lab-Mock-27_S209 | Galveston-Ae-aegypti-Lab-Mock-27_S209_L001_R1_001 (paired).R1.fastq.gz  Galveston-Ae-aegypti-Lab-Mock-27_S209_L001_R1_001 (paired).R2.fastq.gz | Lab | Unexposed | Galveston | Batch_1 |
| Galveston-Ae-aegypti-Lab-Mock-28_S210 | Galveston-Ae-aegypti-Lab-Mock-28_S210_L001_R1_001 (paired).R1.fastq.gz  Galveston-Ae-aegypti-Lab-Mock-28_S210_L001_R1_001 (paired).R2.fastq.gz | Lab | Unexposed | Galveston | Batch_1 |
| Galveston-Ae-aegypti-Lab-Mock-29_S211 | Galveston-Ae-aegypti-Lab-Mock-29_S211_L001_R1_001 (paired).R1.fastq.gz  Galveston-Ae-aegypti-Lab-Mock-29_S211_L001_R1_001 (paired).R2.fastq.gz | Lab | Unexposed | Galveston | Batch_1 |
| Galveston-Ae-aegypti-Lab-Mock-30_S212 | Galveston-Ae-aegypti-Lab-Mock-30_S212_L001_R1_001 (paired).R1.fastq.gz  Galveston-Ae-aegypti-Lab-Mock-30_S212_L001_R1_001 (paired).R2.fastq.gz | Lab | Unexposed | Galveston | Batch_1 |
| Galveston-Ae-aegypti-Lab-Mock-31_S213 | Galveston-Ae-aegypti-Lab-Mock-31_S213_L001_R1_001 (paired).R1.fastq.gz  Galveston-Ae-aegypti-Lab-Mock-31_S213_L001_R1_001 (paired).R2.fastq.gz | Lab | Unexposed | Galveston | Batch_1 |
| Galveston-Ae-aegypti-Lab-Mock-32_S214 | Galveston-Ae-aegypti-Lab-Mock-32_S214_L001_R1_001 (paired).R1.fastq.gz  Galveston-Ae-aegypti-Lab-Mock-32_S214_L001_R1_001 (paired).R2.fastq.gz | Lab | Unexposed | Galveston | Batch_1 |
| Galveston-Ae-aegypti-Lab-Mock-33_S215 | Galveston-Ae-aegypti-Lab-Mock-33_S215_L001_R1_001 (paired).R1.fastq.gz  Galveston-Ae-aegypti-Lab-Mock-33_S215_L001_R1_001 (paired).R2.fastq.gz | Lab | Unexposed | Galveston | Batch_1 |
| Galveston-Ae-aegypti-Lab-Mock-34_S216 | Galveston-Ae-aegypti-Lab-Mock-34_S216_L001_R1_001 (paired).R1.fastq.gz  Galveston-Ae-aegypti-Lab-Mock-34_S216_L001_R1_001 (paired).R2.fastq.gz | Lab | Unexposed | Galveston | Batch_1 |
| Galveston-Ae-aegypti-Lab-Mock-35_S217 | Galveston-Ae-aegypti-Lab-Mock-35_S217_L001_R1_001 (paired).R1.fastq.gz  Galveston-Ae-aegypti-Lab-Mock-35_S217_L001_R1_001 (paired).R2.fastq.gz | Lab | Unexposed | Galveston | Batch_1 |
| Galveston-Ae-aegypti-Lab-Mock-36_S218 | Galveston-Ae-aegypti-Lab-Mock-36_S218_L001_R1_001 (paired).R1.fastq.gz  Galveston-Ae-aegypti-Lab-Mock-36_S218_L001_R1_001 (paired).R2.fastq.gz | Lab | Unexposed | Galveston | Batch_1 |
| Galveston-Ae-aegypti-Lab-Mock-37_S219 | Galveston-Ae-aegypti-Lab-Mock-37_S219_L001_R1_001 (paired).R1.fastq.gz  Galveston-Ae-aegypti-Lab-Mock-37_S219_L001_R1_001 (paired).R2.fastq.gz | Lab | Unexposed | Galveston | Batch_1 |
| Galveston-Ae-aegypti-Lab-Mock-38_S220 | Galveston-Ae-aegypti-Lab-Mock-38_S220_L001_R1_001 (paired).R1.fastq.gz  Galveston-Ae-aegypti-Lab-Mock-38_S220_L001_R1_001 (paired).R2.fastq.gz | Lab | Unexposed | Galveston | Batch_1 |
| Galveston-Ae-aegypti-Lab-Mock-39_S221 | Galveston-Ae-aegypti-Lab-Mock-39_S221_L001_R1_001 (paired).R1.fastq.gz  Galveston-Ae-aegypti-Lab-Mock-39_S221_L001_R1_001 (paired).R2.fastq.gz | Lab | Unexposed | Galveston | Batch_1 |
| Galveston-Ae-aegypti-Lab-Mock-40_S222 | Galveston-Ae-aegypti-Lab-Mock-40_S222_L001_R1_001 (paired).R1.fastq.gz  Galveston-Ae-aegypti-Lab-Mock-40_S222_L001_R1_001 (paired).R2.fastq.gz | Lab | Unexposed | Galveston | Batch_1 |
